# Personalized circuit modeling captures variation in cortical functional connectivity

**DOI:** 10.1101/2024.02.19.581085

**Authors:** Rachel A. Cooper, Murat Demirtaş, Joshua B. Burt, Amber M. Howell, J. Lisa Ji, Grega Repovš, Stamatios N. Sotiropoulos, Alan Anticevic, John D. Murray

## Abstract

Functional magnetic resonance imaging (fMRI) of the human cortex reveals patterns of correlated neural dynamics that are individual-specific and associated with phenotypic variation. However, circuit mechanisms underlying individual variation in functional connectivity (FC) are not well understood. Here, we fit individual-level FC patterns with a biophysically-based circuit model of large-scale cortical dynamics. This model is fit with a small number of neurophysiologically interpretable parameters, and incorporates a hierarchical gradient in local synaptic strengths across cortex parameterized via the structural MRI-derived T1w/T2w map. We applied our modeling framework to resting-state fMRI FC from a large cohort of subjects (N=842) from the Human Connectome Project. We found that the model captures a substantial portion of individual variation in FC, especially with personalized degrees of local synaptic specialization along the hierarchical gradient. Furthermore, the model can capture to the within-subject variation in FC across scans. Empirically, we found that principal modes of individual variation in FC follow interpretable topographic patterns. We developed a framework to assess model expressivity via how these empirical modes of FC variation align with variations in simulated FC induced by parameter perturbations. This framework reveals a straightforward mapping between key parameters and the leading modes of variation across subjects and provides a principled approach to extending computational models. Collectively, our modeling results establish a foundation for personalized computational modeling of functional dynamics in large-scale brain circuits.

## Introduction

The human cortex exhibits substantial variation across individuals, both anatomically and functionally. This variation can be observed macroscopically via magnetic resonance imaging (MRI) and through functional and anatomical relationships with behavioral measures (Finn et al., 2015; Amico and Goñi, 2018; Mueller et al., 2013; Smith et al., 2015; Forstmann et al., 2010; Cohen et al., 2009; Saygin et al., 2012, 2016; Chowdhury et al., 2013). However, the neural circuit mechanisms of functional variation remain unclear. Two elements which in combination potentially make substantial contributions to this across-subject variation are long-range white-matter anatomical connectivity and cortical physiological dynamics. However, the extent of the contributions of each remains unknown.

Empirical studies have shown that subject-level cortical functional connectivity (FC) patterns, as measured by resting-state functional MRI (fMRI), contains meaningful information about individual humans. The functional connectome is subject-specific and stable over time — at least on the timescale of months — and across resting-task conditions (Finn et al., 2015; Amico and Goñi, 2018; Laumann et al., 2015; Gordon et al., 2017; Miranda-Dominguez et al., 2014; Demeter et al., 2020; Gratton et al., 2018). FC features are predictive of individual differences in behavioral and cognitive measures (Finn et al., 2015; Amico and Goñi, 2018; Mueller et al., 2013; Smith et al., 2015). FC shares many properties with anatomical connectivity, as measured through diffusion-weighted imaging tractography; both functional and anatomical connectivity exhibit stronger across-subject similarity with genetic relatedness (Demeter et al., 2020; Smith et al., 2021). Both functional and anatomical connectivity variation across subjects are heterogeneous across the cortex, with greater differences in higher-order association areas than in sensorimotor areas (Finn et al., 2015; Mueller et al., 2013; Laumann et al., 2015; Miranda-Dominguez et al., 2014; Demeter et al., 2020; Gratton et al., 2018; Sun et al., 2022; Xu et al., 2019; Kong et al., 2018; Hill et al., 2010; Sun et al., 2022). Within-subject differences follow a contrasting gradient, with greater variation found in sensorimotor regions (Laumann et al., 2015; Gratton et al., 2018; Sun et al., 2022; Kong et al., 2018). Further, across both time and task conditions, the magnitude of across-subject variation is much larger than within-subject variation (Laumann et al., 2015; Gratton et al., 2018). However, anatomical connectivity is only weakly predictive of subject-specific functional connectivity (Zimmermann et al., 2018a; Messé, 2020), which may be due in part to limitations with current estimates of single-subject level anatomical connectivity from diffusion tractography.

An approach to elucidate the mechanistic origins of functional variation across individuals is to use biophysically-based neural circuit models of resting-state functional dynamics. One such model construction simulates the cortex as a network of coupled excitatory and inhibitory units, analogous to populations of excitatory pyramidal neurons and inhibitory interneurons (Deco et al., 2014; Demirtaş et al., 2019). This approach bridges multiple spatial scales, allowing macroscopically observable large-scale brain dynamics to be simulated from cellular-level components. The units interact via recurrent local interactions as well as long-distance excitatory connections. The free parameters are directly related to either synaptic cortical dynamics or anatomical connectivity and can be quantitatively fit to empirical FC patterns. This modeling framework thereby provides a useful testbed for evaluating the relationships between functional connectivity, anatomical connectivity, and cortical activity dynamics. (Demirtaş et al., 2019) expanded this model to incorporate regional variation in the strengths of local recurrent synaptic connections, following a hierarchical gradient indexed by the the T1-to T2-weighted (T1w/T2w) structural MRI measure, based on convergent findings of microcircuit specialization (Burt et al., 2018). However, it remains unclear if these models are sensitive enough to capture differences in FC across individuals.

Here, we modeled functional connectivity at the individual level for subjects from the Human Connectome Project (Essen et al., 2013), using the aforementioned circuit model of (Demirtaş et al., 2019). Optimizing subject-level synaptic parameters yielded a model capable of capturing individual-level FC patterns with subject specificity. Delving deeper, we decomposed the empirical FC patterns into principal components of variation across subjects. We found that the components corresponding to the greatest across-individual variation are highly correlated with neurobiologically interpretable neural features. We further developed a sensitivity analysis method of quantifying a model’s ability to capture across-subject differences, which can guide parametric extensions of the model.We found that subject-level structural connectivity consistently failed to improve the subject-level simulated FC. As proof-of-concept, we examined within-subject variability across scans, finding subject-specific stability over time empirically, which we reproduced with the model using personalized synaptic parameters. All together, our model supports the hypothesis that microcircuit properties related to cortical physiology and dynamics make dominant contributions to functional variation across individuals in healthy populations.

## Results

### Personalized circuit modeling framework

Our framework for personalized circuit modeling of large-scale cortical dynamics builds upon a model with a small number of free fitting parameters which are neurophysiologically interpretable (Demirtaş et al., 2019). We fit the model at the individual-subject level to the cortical functional connectivity (FC) matrices of 842 subjects from the the Human Connectome Project (HCP) (Essen et al., 2013), using the HCP-MMP parcellation with 180 areas per hemisphere (Glasser et al., 2016). The cortex is modeled as a network of pairwise-connected nodes, where each parcel is represented as a node. The activity within each parcel is approximated by a circuit consisting of one excitatory unit (*E*) and one inhibitory unit (*I*) (Fig. 1A). The excitatory and inhibitory units interact locally (represented by the parameters 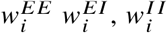 and 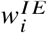). The excitatory units also receive long-range inputs from other nodes whose strengths are set the structural connectivity (SC) between parcels *i* and *j*, which is scaled by the free fitting parameter for global coupling, *G*. The SC matrix is derived via diffusion tractography from the HCP dataset (Demirtaş et al., 2019). Here we primarily use a group-level average SC (from a subset of 328 subjects in the HCP dataset), to isolate the ability of the neurophysiologically interpretable synaptic parameters to capture individual variation in FC (Fig. 1B).

**Figure 1.**
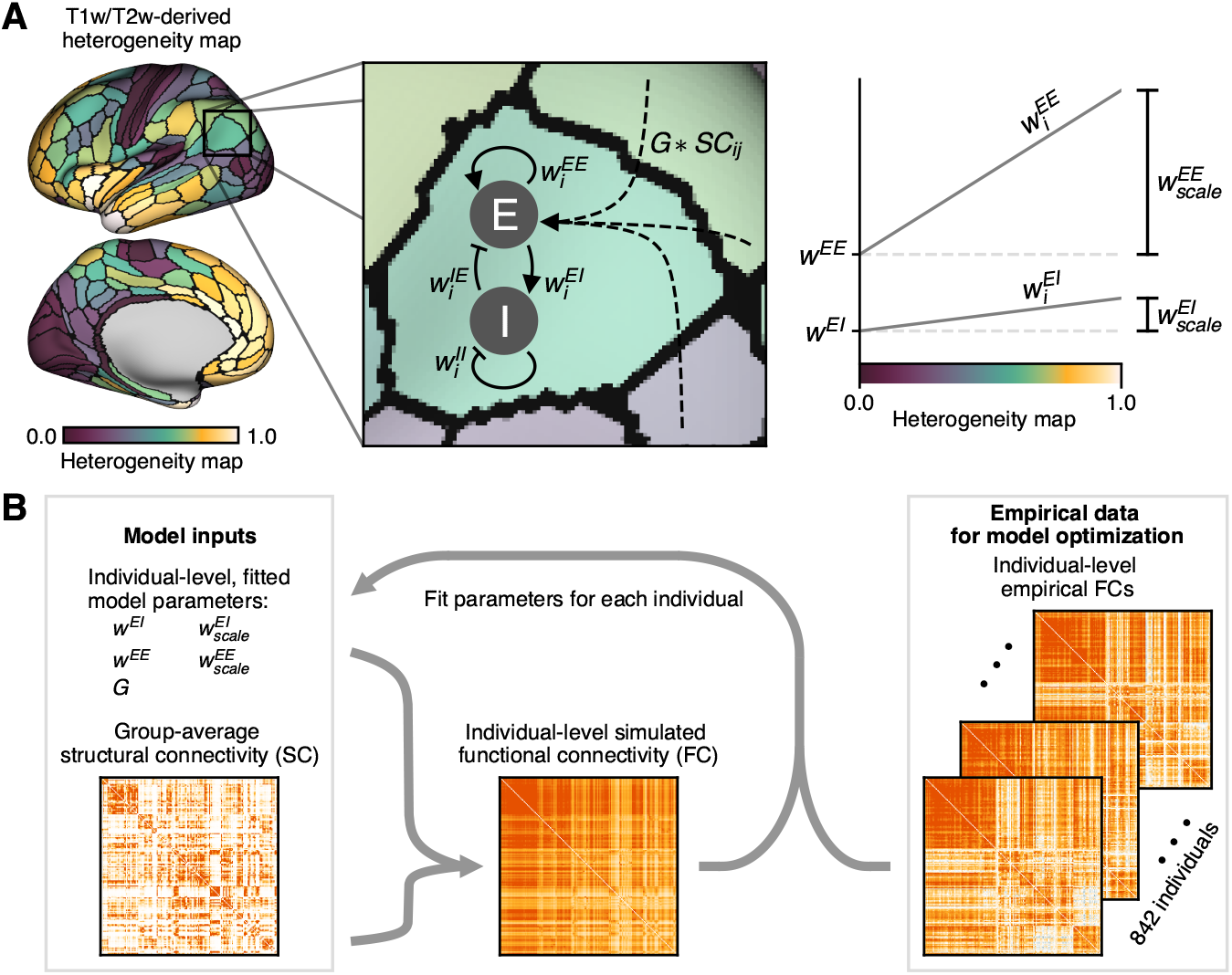
Schematic depicting the circuit model simulation of individual-level FC matrices. **(A)** Schematic of the heterogeneous circuit model. Each parcel is modeled as a circuit containing a single excitatory (*E*) unit and a single inhibitory (*I*) unit. The *I* unit inhibits itself recurrently, as well as the local *E* unit. The *E* unit excites itself recurrently, the local *I* unit, and the *E* units in every other parcel, scaled according to the global coupling parameter,*G*, and the structural connectivity (SC) between parcels *i* and *j*. The weights 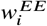 and 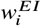 are scaled across cortical areas according to a hierarchical gradient based on linearized T1w/T2w maps, meaning units in higher-order association areas generally have higher local recurrent weights than those in sensorimotor areas. **(B)**Schematic of the modeling flow. Given any vector of model parameter values (chosen here using a stochastic optimization algorithm) and an SC matrix (here, the group-level SC matrix), functional connectivity (FC) matrices were generated using the circuit model depicted in panel **(A)**. Individual-level parameter values are iteratively, semi-stochastically tuned to maximize the fit to their respective empirical FC matrices. The heterogeneous model, shown in panel **(A)**, has five free parameters to fit: 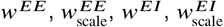, and *G*.

Our circuit model is described as heterogeneous, in that local excitatory synaptic strengths vary across cortical areas following a sensory-association hierarchical gradient, parametrized by the T1-weighted/T2-weighted (T1w/T2w) maps as a proxy for cortical hierarchy (Fig. 1A left and right) (Burt et al., 2018; Demirtaş et al., 2019). The magnitude of dependence of 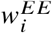 and 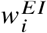 thereby depend on a T1w/T2w-derived heterogeneity map with intercepts *w*^*EE*^ and *w*^*EI*^ and slopes 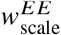and 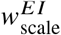, respectively (Fig. 1A right). A homogeneous model, with uniform local excitatory strengths across areas, sets slopes 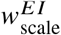 and 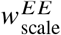 to zero. The parameters 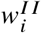 and 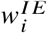 are then fixed to maintain a uniform baseline firing rate of ∼3 Hz. These five free parameters — 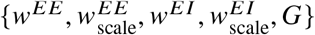 — are fit using empirical data to return the optimal simulated FC matrix (Fig. 1B). As examples, the empirical and simulated FC matrices for two individuals and the group-average visually demonstrate the model’s ability to produce distinguishable FC matrices with only synaptic parameters varying across individuals (Fig. 2A).

**Figure 2.**
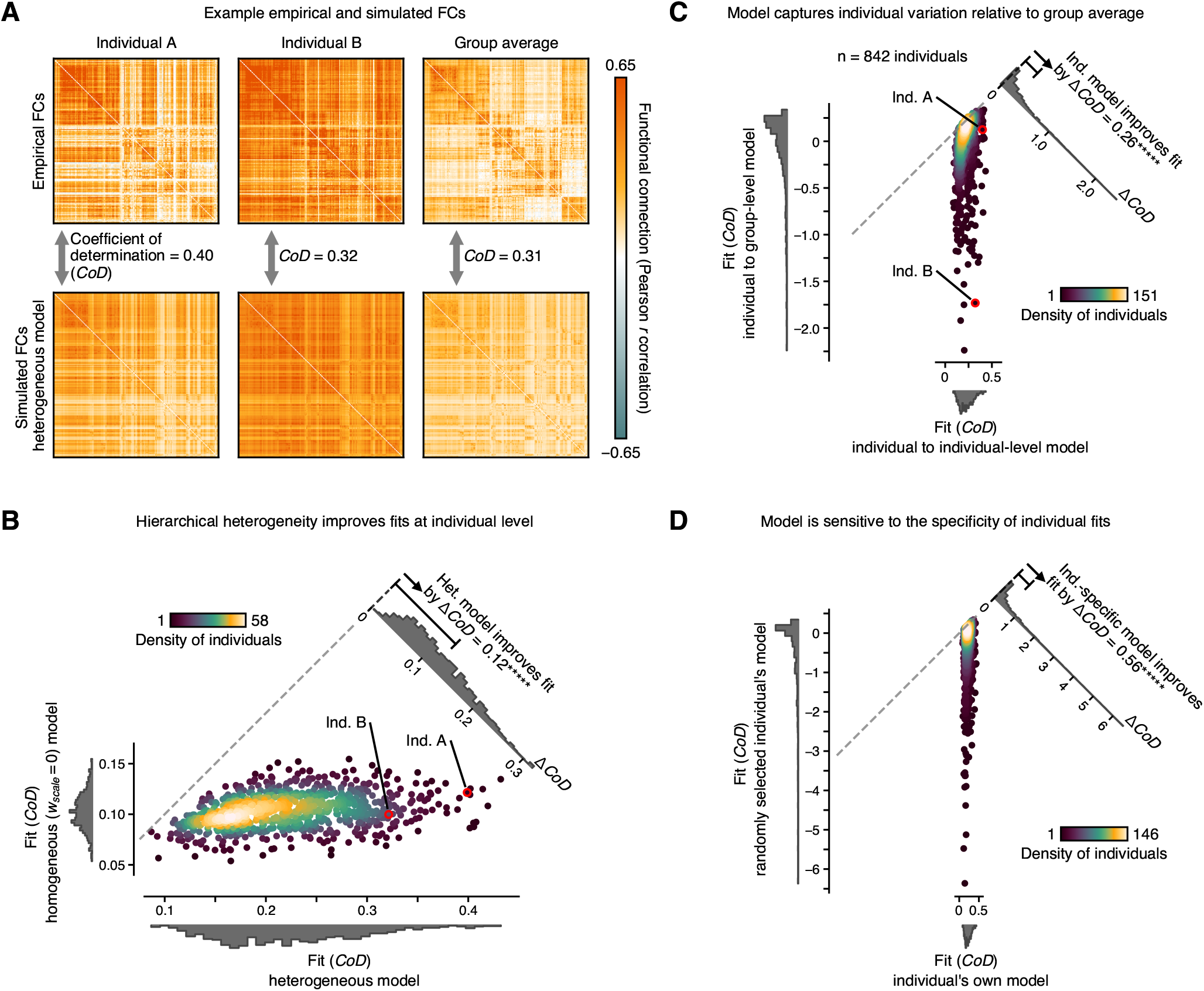
Heterogeneous cortical circuit model is capable of capturing individual-level variation by tuning synaptic dynamics parameters. **(A)** Empirical and simulated FC matrices for two subjects and the group average. CoD refers to the coefficient of determination, a similarity measure which is similar to but more strict than the square of the Pearson correlation coefficient (Fig. S1A–E). The simulated FC matrices visually differ from each other and capture many of the patterns visible in the respective empirical FC matrices via optimizing synaptic parameters. **(B)** Heterogeneity across cortex is important for individual-level modeling. Individual-level, simulated FC matrices were generated for each of the heterogeneous and homogeneous models. In the homogeneous model, local circuit parameters 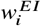 and 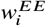 are constant for all parcels across the entire cortex, which is algorithmically equivalent to the heterogeneous model with 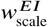 and 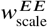 clamped to zero. Incorporating heterogeneity across the cortex into the model according to corticalhierarchygreatly improves fits, particularly for subjects well-fit by the homogeneous model. Each point in the scatterplot corresponds to an individual. Fits are measured via CoD values. Color/brightness of the scatter points indicate the relative density of scatter points in that region (*****, *p <* 10^−5^, bootstrapping across individuals). **(C)** Individual-level functional differences are captured with personalized synaptic parameters. Each subject was fit to a simulated group-level FC matrix (y-axis) and the subject’s own simulated individual-level FC matrix (x-axis). Subject-level fits are consistently improved with the individual-level model, with nearly forty percent of subjects having fit values less than zero relative to the group-level model, meaning the group-level model is insufficient for subject-level modeling. There is a high concentration of subjects clustering near the unity line with a long tail favoring the subject-level simulated FC matrices. This indicates that while some subjects were fit reasonably well by the group-level synaptic parameters, many others were fit very poorly (*****, *p <* 10^−5^, bootstrapping across individuals). **(D)** Synaptic parameters produce simulated FC matrices that are subject-specific. Subjects were fit to a randomly selected individual’s simulated FC matrix (y-axis) as well as their own simulated FC matrix (x-axis). Approximately half of the subjects have a fit value less than zero relative to the randomly-selected subject’s simulated FC matrix, implying that the synaptic parameters for one subject often fail to predict the functional activity of another subject. Again, subjects cluster near the unity line with a long tail favoring the within-subject simulated FC matrices, indicating that some subjects were fit reasonably well by the across-individual synaptic parameters, but many others were fit very poorly (*****, *p <* 10^−5^, bootstrapping across individuals).

The choice of fitting metric used to quantify the similarity between the simulated and empirical FC matrices inherently influences characteristics of the optimal simulated FC matrices. For example, Pearson correlation coefficient is insensitive to differences in slope and intercept (Fig. S1A–E), meaning it is blind to these biologically meaningful structures. As suggested by Scheinost et al. (2019), we chose to primarily perform our fitting procedures and subsequent analysis using “prediction *r*^2^” (see Methods), which we generically denoted as the “coefficient of determination” (CoD) throughout this study. For highly correlated sets of data, CoD values converge to the square of the Pearson correlation coefficient.

### Synaptic parameters capture individual differences in functional connectivity

Most previous large-scale modeling studies assume homogeneous distributions of local circuit parameters across the cortex (Zimmermann et al., 2018b; Aerts et al., 2018, 2020; Deco et al., 2013, 2014; Schirner et al., 2018). A modeling approach allowing local circuit parameters to vary across cortex was implemented by Demirtaş et al. (2019), which found that heterogeneity in local circuit properties following a hierarchical gradient substantially improves the model fit to empirical FC, relative to a homogeneous model, at the group level. Similar hierarchical structure has emerged previously from more generalized models of FC (Wang et al., 2019; Singh et al., 2020; Kong et al., 2021). At the single-subject level, we similarly found that the model-empirical fit values of the heterogeneous model are greatly improved relative to a homogeneous model (Fig. 2B). In these scatter plots, each point represents an individual subject, and the brightness/color depicts the relative density of scatter points on the plot. Scatter points close to the dashed, unity line represent subjects who are fit similarly well by either model.

Incorporating hierarchical heterogeneity into the synaptic parameters of the model approximately doubles subjects’ fit values (Δ*CoD* = 0.12 on average). Interestingly, subjects well-fit by a homogeneous model were disproportionately improved by the introduction of heterogeneity (Fig. 2B). Hierarchical heterogeneity is evident in the simulated FC matrices themselves; at both the individual and group levels, the difference between the simulated heterogenous and homogeneous FC matrices clearly exhibit hierarchical structure (Fig. S2). At the individual level, the heterogeneous model is likewise able to capture the hierarchical topography of dissimilarity across subjects better than the homogeneous model. Notably, the model with hierarchical heterogeneity remains low-dimensional, adding only two new free parameters, and these parameters are generally not redundant with each other (Fig. S3).

Personalizing the synaptic parameters (without personalizing SC) enables individualized FC patterns to be captured by the simulated FC matrices. This is true relative to the group-average simulated FC matrix (Fig. 2C) as well as the individual-level simulated FC matrix of a randomly selected subject (Fig. 2D). Thus, the simulated FC matrices show individual subject specificity.

Some subjects were fit reasonably well by the group and/or across-individual synaptic parameters, but many others were fit very poorly. Values of the similarity metric, CoD, fall within the range (-∞, 1], where CoD = 0 signifies a model that does not explain any variance in empirical data. Thus, the individualized simulated FC matrices embody a substantial ability to capture individual characteristics (ΔCoD =0.26, better than the group-average model on average and ΔCoD =0.56, better than the across-subject model on average). Further, with model-empirical fit values less than zero, over half of the subjects were not captured in any meaningful way by the group or across-subject simulated FC (Figs. 2C,D). These effects are not attributable to overfitting. There are 16,110 unique off-diagonal edges in a 180 ×180 FC matrix fit with 5 free parameters. We confirmed minimal impact of overfitting by splitting empirical time-series data for training and testing and comparing testing data to the model fit to training data (Fig. S1F–H).

### Individual variation in functional connectivity has biologically interpretable structure

What are the dominant empirical modes of functional variation across subjects? To characterize this, we applied principal component analysis (PCA) to the set of 842 FC matrices (Fig. 3A). Visualizing the three leading principal components (PCs) reveals patterns that may be informative as to neurobiological contributions to individual differences in FC (Fig. 3B–D). We can also visualize, for each PC matrix, its leading component which is a spatial map.

**Figure 3.**
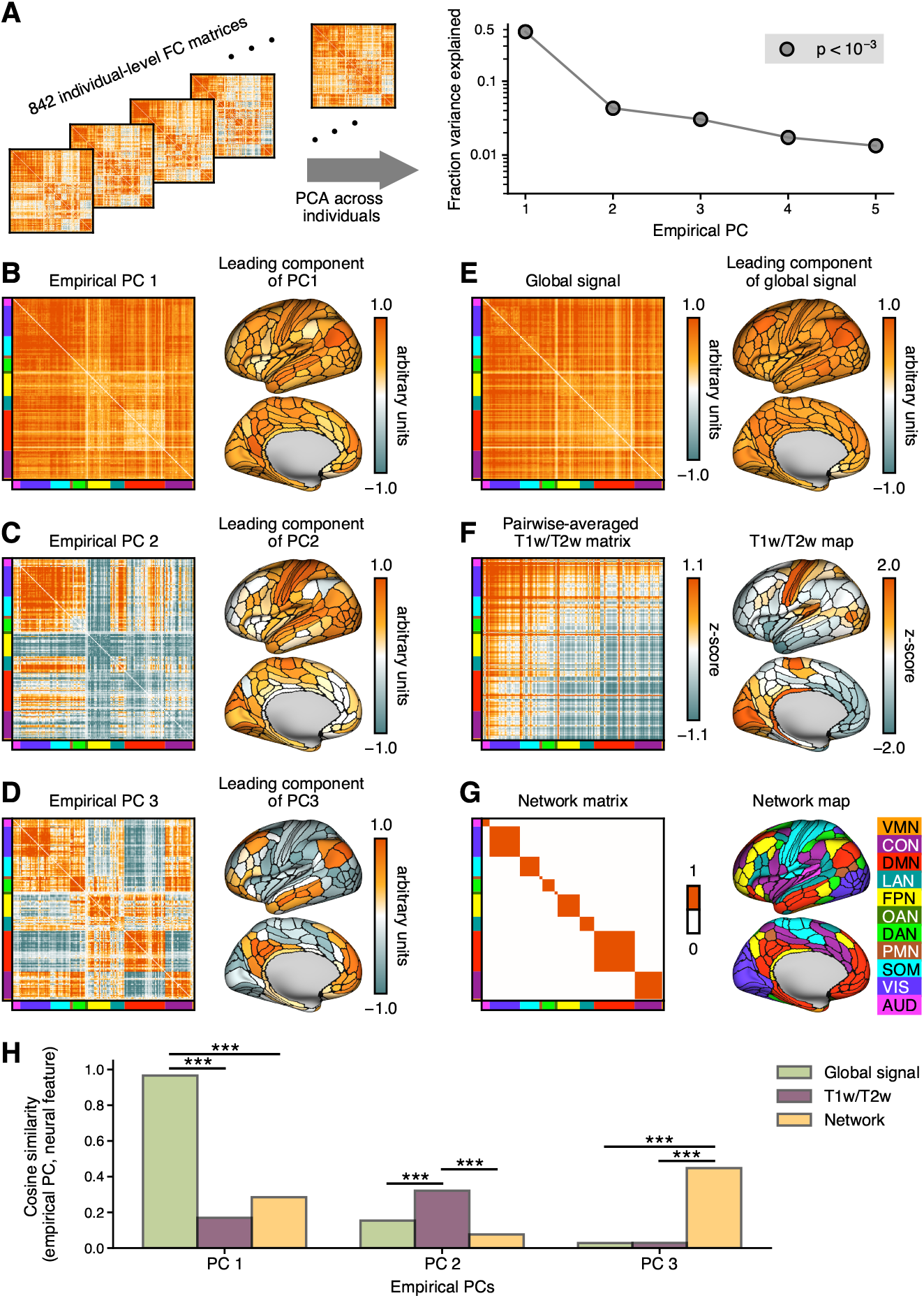
Empirical across-individual variation is correlated with neural features. **(A)** Schematic of performing principal component analysis (PCA) across individuals and the resulting scree plot for the leading 5 principal components. PCA was performed across individuals, yielding an ordering of principal component matrices, which represent the axes of greatest across-subject variation. As evidenced by the scree plot, the first three principal components capture over fifty percent of across-individual variation, which is highly significant over the null hypothesis that all across-individual variation is an artifact of the finite timescales of fMRI scans. Notably, our data was not processed with global signal regression (GSR), which may explain the large amount of variance explained by the first principal component. **(B, C, D, left)** First, second, and third principal components of the empirical FC matrices taken across subjects, showing the three axes of greatest variation across individuals in the empirical data. **(B, C, D, right)** Shown in the brain maps, the leading component of variation across brain areas of each of the respective PC matrices mapped onto the brain. **(E)** Global signal matrix, calculated as the difference between the empirical FC matrices with and without GSR applied, averaged over individuals, and its axis of greatest spatial variation mapped onto the brain. This matrix is highly correlated with the first empirical PC (cosine similarity = 0.97). **(F)** Hierarchy matrix, where each element *T*_*i j*_ represents the average T1w/T2w value of parcels and (left), and the group-average T1w/T2w map plotted on the brain (right). The hierarchy matrix is highly correlated with the second empirical PC (cosine similarity = 0.32). **(G)** Network matrix and map, denoting parcels with similar functional characteristics, as determined by Ji et al. (2019). The network matrix is highly correlated with the third empirical PC (cosine similarity = 0.45). The networks are defined as follows: VMN: ventral multimodal, CON: cingulo-opercular, DMN: default mode, LAN: language, FPN: frontoparietal, OAN: orbito-affective, DAN: dorsal attention, PMN: posterior multimodal, SOM: somatomotor, VIS: visual, AUD: auditory. **(H)** Cosine similarity between leading three principal components and the neural features shown in E, F, & G. PC1 is highly correlated with global signal, significantly more than both T1w/T2w and network structure. PC2 is highly correlated with hierarchy (significantly more than both global signal and network structure), and PC3 is highly correlated with the network structure (significantly more than both global signal and hierarchy) (***, *p <* 10^−3^, calculated by bootstrapping across individuals). An analogous plot using Pearson correlation coefficient as the similarity metric is shown in Fig. S4A.

The first principal component, PC1, exhibits a pattern with the same sign across edges (Figs. 3B), which we hypothesized may relate to capturing individual differences in global signal. Global signal is known to vary substantially across individuals, time, and conditions (Li et al., 2019; Orban et al., 2020; Yang et al., 2014). To derive an edge-level matrix corresponding to global signal, we defined the global signal matrix as the difference between the group-average empirical FC matrices with and without global signal regression (GSR) applied (Fig. 3E). We found that principal component 1 (PC1, Fig. 3B) is highly correlated with this estimate of global signal (cosine similarity = 0.97). To evaluate the robustness of this correlation, we also partition the subjects into subgroups of 40 and calculate the principal components of variation across subjects for each subgroup. These subgroup PC1s and the global signal matrix are also strongly correlated (cosine similarity = 0.95 ±0.01).

PC2 has a pattern resembling a hierarchical gradient, with sensorimotor-sensorimotor edges having generally positive values, and association-association edges having generally negative values (Fig. 3C). We defined the T1w/T2w matrix such that each element *T* is the average T1w/T2w value of parcels *i* and *j* (Fig. 3F). We found a moderately high level of similarity between the T1w/T2w matrix and PC2 (cosine similarity = 0.32; subgroup cosine similarity = 0.25 ±0.07). This prominence of a hierarchical pattern clarifies by adding hierarchical heterogeneity as parameters in the model substantially improves model-empirical fit (Fig. 2B), and justifies that model extension as a useful approach to capture individual differences in FC.

PC3 of the empirical FC matrices shows a distinctive pattern resembling commonly defined functional network assignments, with same-network edges tending to have greater values than across-network pairs (Fig. 3D). We defined the binary network map as having a value of one for same-network pairs of parcels and a value of zero for across-network pairs, based on the functional network mapping defined by Ji et al. (2019) (Fig. 3G). The network map and PC3 are correlated (cosine similarity = 0.45; subgroup cosine similarity = 0.30 ± 0.09), which suggests that network structure likely plays an important role in across-subject variation. This will be explored in subsection 2.5 to guide further model extensions. Figure 3H and Supplementary Figure S4A summarize the correlations between the PCs and each of the neural features using cosine similarity and Pearson correlation coefficient as similarity metrics, respectively.

### Heterogeneous model captures leading modes of individual variation

Individualized synaptic model parameters are generally sufficient to capture subject-level FC patterns and distinguish between subjects by a large margin (Fig. 2). We explored how expressivity of a model, for a given parameterization, relates with the modes of variation across subjects as represented by the leading principal components of empirical across-individual variation (Fig. 3). To that end, we developed a method of quantifying the relationships between leading empirical PCs across subjects and the simulated FC matrices.

This method for any simulated FC is as follows. The set optimal synaptic parameter values, 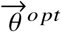, were found (Fig. 4A, left box; 5-parameter heterogeneous model shown). One of these optimal values was altered by a small amount, resulting in a set of perturbed parameters 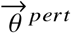. These perturbed parameter values were used to simulate the perturbed FC matrix *FC* 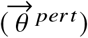, demon-strating the effect of altering one of the parameter values on the simulated FC. We defined and calculated the *0* sensitivity matrix as the difference between the optimal and perturbed simulated FC matrices scaled by the magnitude of the perturbation (for example, Fig. S4B). This was repeated for each free parameter, yielding one sensitivity matrix per free parameter (Fig. 4A, center boxes). The linear combination of these sensitivity matrices defines a subspace of FC-space that locally approximates what FC patterns the model can generate under this parameterization. A PC can then be projected onto this sensitivity subspace, to calculate its overlap or alignment with the subspace, thereby quantifying the model’s ability to capture the mode of FC variation described by that PC (Fig. 4A, right).

**Figure 4.**
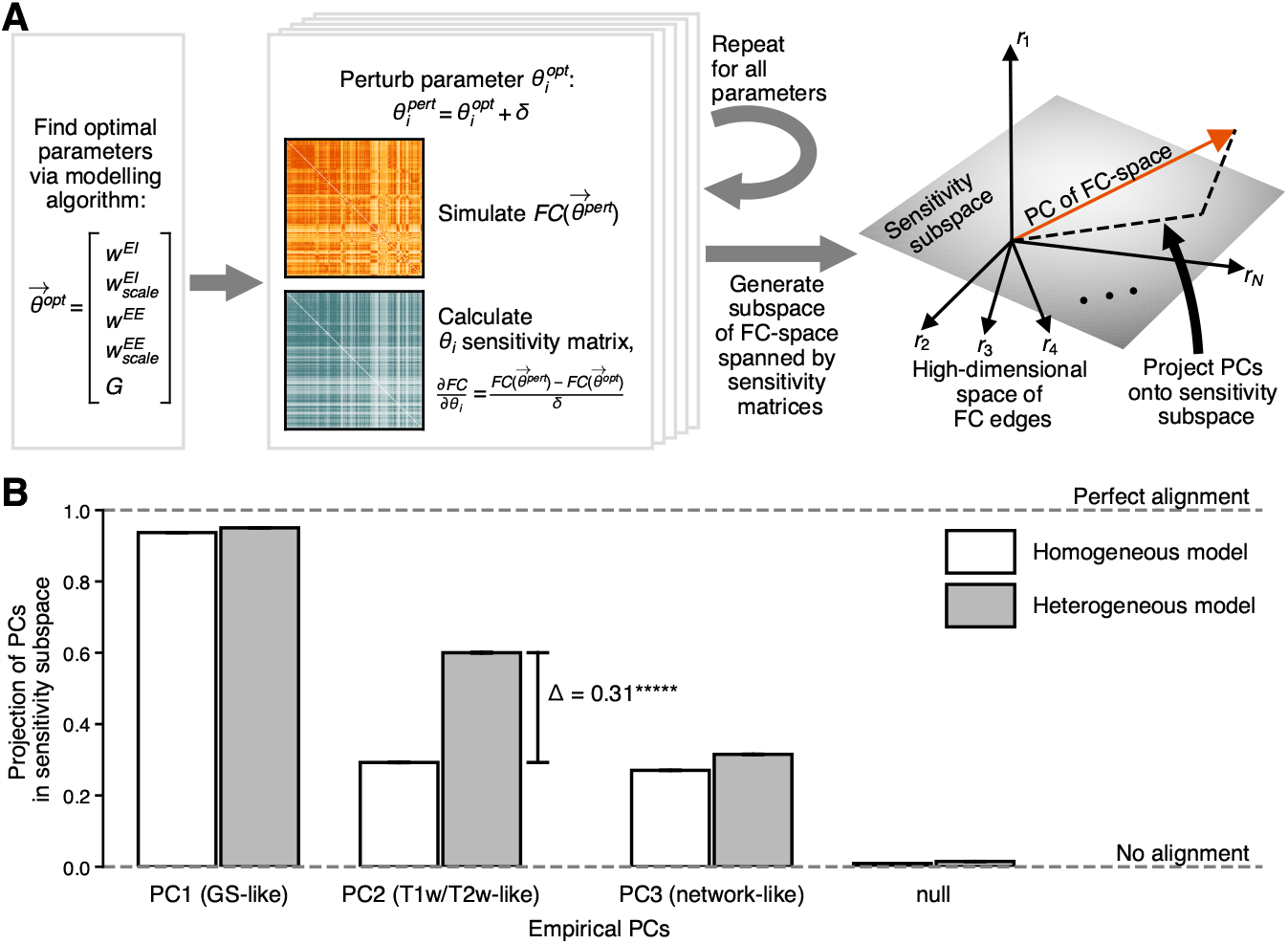
A hierarchy-based distribution of local circuit parameters across the cortex improves the model’s sensitivity to across-subject variation. **(A)** Schematic illustrating the projection of leading empirical principal components onto the subspace of FC-space to which the model is sensitive. Using the subject-level sets of optimal modeling parameters 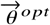 (generated by the modeling algorithm), one of the optimal parameters was perturbed by increasing or decreasing its value slightly. The set of parameters termed 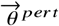 refers to the set of optimal parameters with one value replaced by its perturbed value. We simulated the FC matrix corresponding to the parameters 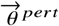. We defined the *θ*_*i*_ sensitivity matrix as the difference between the optimal and perturbed simulated FC matrices divided by the difference between the optimal and perturbed parameter *θ*_*i*_. This can also be written as 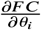. This is repeated for each parameter, generating five (heterogeneous model) or three (homogeneous model) sensitivity matrices. Flattening the matrices into vectors and taking their linear combination generates a low-dimensional subspace of FC-space which locally approximates the region of FC-space accessible by the modeling parameters. We then projected the leading empirical PCs onto this subspace, finding the alignment (or fractional projection) of the PCs onto the sensitivity subspace. This alignment represents an estimate of the model’s ability to capture that mode of variation. **(B)** Alignment of the leading empirical PCs onto the sensitivity subspaces defined by the homogeneous and heterogeneous models. Both models capture PC1 similarly well and PC3 similarly poorly, with the heterogeneous model generating slight but statistically significant improvements to both. The heterogeneous model, however, is considerably more capable of capturing PC2 for a majority of subjects (*****, *p <* 10^−5^, bootstrapping across individuals). The mean fractional projections of PC1 and PC3 are also improved, albeit modestly (/1 = 0.013, p < 10^−5^ and Δ = 0.045, p < 10^−5^, respectively). The standard error of the mean for each bar is depicted with an error bar, all of which are nearly too small to be distinguished.

We projected the leading empirical across-subject PCs (Figs. 3B–D) onto the sensitivity sub-space of each individual’s fitted models: the 5-dimensional heterogeneous model, and the 3-dimensional homogeneous (i.e., 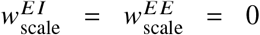) model (Fig. 4B). We found that the first PC is captured well by both models. Since PC1 is highly correlated with our estimate of global signal (Figs. 3B,E,H) and global signal is responsible for much of the structure of the simulated FC matrices (Aquino et al., 2022), it is reasonable for homogeneous and heterogeneous models to capture PC1 similarly well. However, global signal varies also greatly within-subject S5; modeling individual variation thus requires higher-order PCs to be captured. By incorporating hierarchical heterogeneity across the cortex into the model, the mean fractional projection of PC2 is improved substantially (Δ = 0.31). However, the third PC is captured similarly poorly by both models.

The strong alignment between the leading PCs and the sensitivity subspaces is especially striking given that these subspaces are 3- or 5-dimensional (homogeneous or heterogeneous model), while FC-space is 16,110-dimensional (for each of the unique off-diagonal elements of a 180 × 180 FC matrix). This is reflected by the low alignment of a randomly-generated null vector with the model sensitivity spaces. As a proof-of-concept demonstration that this novel method of quantifying feature alignment imparts information about the underlying model, we also projected various neural features (Figs. 3E–G) onto the subject-level sensitivity subspaces (Fig. S4C). As expected, the models with a hierarchically-defined heterogeneous distribution of synaptic parameters were more capable of capturing the T1w/T2w matrix than the model with a homogeneous distribution. This suggests that the alignments being reported from this method accurately represent the structure of the respective model.

Results above (Figs. 2B and 3C,F) show that a heterogeneous cortical model improves fit values because of the strong alignment between the leading components of variation across individuals and hierarchy (which is used here to incorporate heterogeneity into the model). Our analysis combining PCA with the subject-level models supports this hypothesis. These findings suggests that the physiological regime of neural dynamics varies across the cortex, and the degree of the variation across the cortex differs across individuals.

### Within-network correlated noise captures a leading mode of individual variation

The third leading principal component of individual variation, PC3, exhibits block-like structure resembling functional networks (Fig. 3D,F,H). Thus, we tested extending the model by incorporating within-network structure in a variety of ways to identify possible physiological mechanisms that may contribute to variation across subjects. Specifically, we tested adding correlated background noise inputs, network-structured global coupling, and feedforward inhibition (Figs. 5, S6, S7, S8).

**Figure 5.**
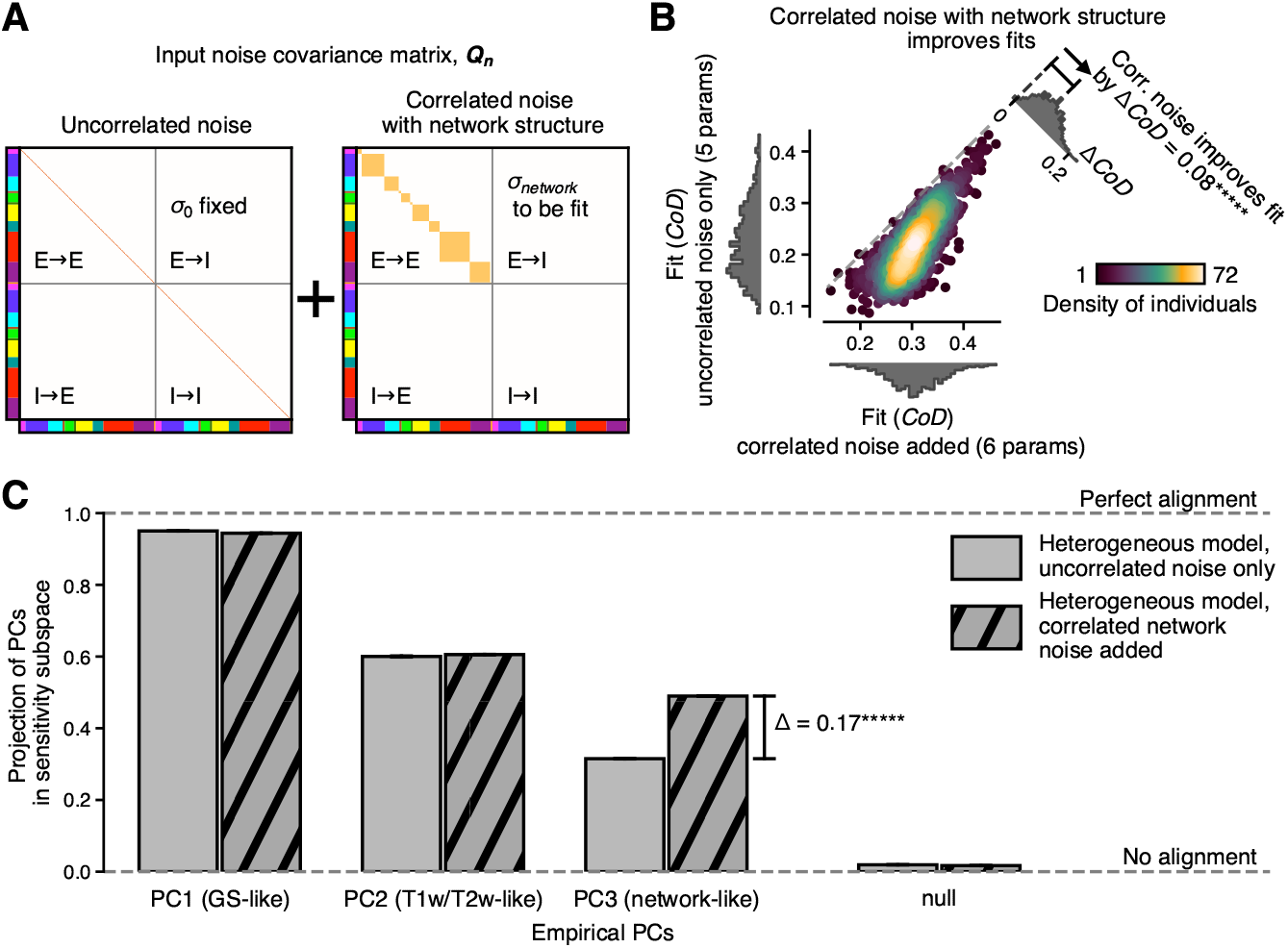
Allowing correlated noise processes between same-network parcels improves subject-level modeling, capturing a leading component of across-individual variation. **(A)** Uncorrelated noise across parcels produces a noise covariance matrix identical to the identity matrix (left panel). We allowed noise between excitatory-to-excitatory units within a common network to be correlated, thus adding a block diagonal component scaled by the new parameter *σ*_network_ to the noise covariance matrix. **(B)** Adding within-network noise scaled by *σ*_network_ significantly improves the model’s ability to replicate subject-level FC matrices, especially for subjects who were not able to be fit well with the heterogeneous model with uncorrelated noise (*****, *p <* 10^−5^, bootstrapping across individuals). **(C)** By projecting the leading empirical PCs onto the sensitivity subspaces defined by the heterogeneous models with and without correlated, within-network noise, we found that the model with correlated noise is substantially better able to capture the third PC.

Among the different model extensions tested, same-network parcels having correlated background noise improved the model fits the most (Figs. 5, S6, S7, S8). In the model, each cortical parcel receives time-varying background noise input. In the preceding sections, the model assumes each neural unit receives independent noise. Thus, the noise covariance matrix **Q**_**n**_, quantifying the covariance of the noise across neural units, is the identity matrix in this case (Fig. 5A, left). To incorporate network structure, we assumed patterned covariance of noise input to same-network excitatory units. This adds a block diagonal structure to the noise covariance matrix (Fig. 5A, right). A sixth parameter, *σ*_network_, is added to the model to scale the strength of the correlation between noise processes.

We modeled each subject with this enhanced model, finding that this extension significantly improves the fit values between the empirical and simulated FC matrices (ΔCoD = 0.08, Fig. 5B). This enhanced model disproportionately augments the fit values for subjects poorly fit by the uncorrelated-noise only model, which is the opposite trend seen from adding heterogeneity to the model (Fig. 2B). Thus, incorporating correlated noise into the model may prove to be especially beneficial to subjects for whom the addition of heterogeneity yielded little improvement. The addition of within-network noise is readily apparent in the simulated FC matrices; at both the individual and group levels, the difference between the heterogeneous modeled FC with and without additional noise show clear network-like structure (Fig. S2). This improved model captures dissimilarity across subjects similarly to the heterogeneous model without network noise, and the additional free parameter is not redundant with the other parameters (Fig. S3).

We performed the previously described sensitivity subspace alignment analysis (Fig. 4A), which specifically provides information about whether extensions to the model are useful for capturing differences across subjects. Using this analysis, we again calculated the projections of the leading three empirical PCs onto the five- and six-dimensional sensitivity subspaces for heterogeneous models without and with correlated same-network noise, respectively (Fig. 5C). The addition of correlated noise markedly improves the model’s ability to capture PC3 (Δ = 0.17). The alignments of the first and second PCs were only slightly altered by this model (Δ = -0.0094 for PC1; Δ = 0.0024 for PC2). This suggests that the improvements in fit achieved from the addition of the *σ* _network_ parameter are specifically useful for capturing variation across individuals along the PC3 mode. We also projected various neural features (Figs. 3E–G) onto the sensitivity matrices of the extended model (Fig. S4C). The network matrix aligned very closely with the sensitivity matrices for the model with network-shaped correlated noise.

### Subject-specific SC fails to improve model

Structural connectivity (SC) matrices, derived from diffusion tractography, are used by the model to represent the strengths of long-range projections between cortical areas. All preceding results used a group-average SC matrix. It is unclear to what extent individual differences in FC can be explained by individual differences in SC. SC connection strengths can vary widely across subjects (Figs. S9A,B), likely due to random noise induced by limitations in tractography (Maier-Hein et al., 2017; Jones et al., 2013; Jeurissen et al., 2017; Sotiropoulos and Zalesky, 2019; Zalesky et al., 2016). However, subject specificity has been observed within FC-SC correlations for some cortical parcellations (Zimmermann et al., 2018a; Messé, 2020), and structural and functional connectomes vary across individuals with similar patterns (Figs. S9C,D and S3A–H).

To examine the utility of modeling with personalized SC, we generated subject-level simulated FC matrices constrained by subject-level SC matrices (Fig. 6A). We found that modeling a subject’s FC matrix using their own subject-level SC matrix failed to produce greater fit values, compared to a model using a randomly selected other subject’s SC matrix (Fig. 6B). Subjects’ FC modeled using the group-average SC matrix were fit significantly, albeit modestly, better than using the individual-specific SC matrices (Fig. 6C). These results hold for models with and without correlated noise added (Fig. S9E,F).

**Figure 6.**
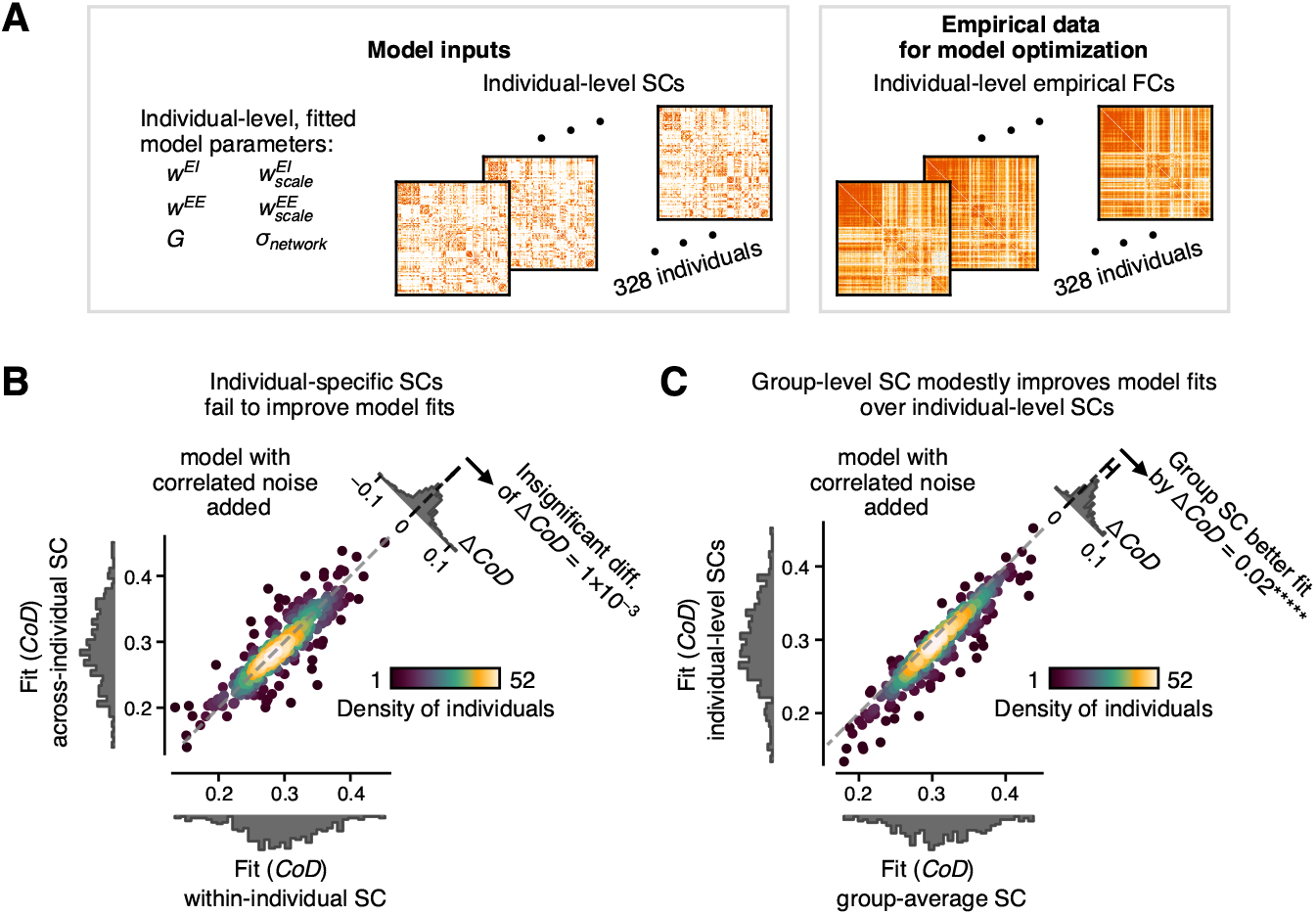
Individual-level heterogeneous model not improved by subject-specific SC matrices, preferring instead the group-averaged SC matrix. **(A)** Schematic of model inputs. We generated simulated subject-level FC matrices by constraining the model with subject-level SC matrices, within and across individuals. **(B)** Model is insensitive to subject-level information in SC matrices. Slightly over half of the subjects (57%) are better fit by the model using their own SC matrix than a randomly chosen subject’s SC matrix. This holds for models with correlated noise (shown here) and those without (Fig. S9) (*p >* 0.1, bootstrapping across individuals). **(C)** Model shows slight preference for group-average SC over individual-level SC matrices. All subjects are fit similarly or better by a model using the group-averaged SC matrix than the subject-level SC matrices. This also holds for models with correlated noise (shown here) and those without (see Figure S9) (*****, *p <* 10^−5^, bootstrapping across individuals).

Importantly, while there was no significant difference in fit values between models seeded using a subject’s own SC matrix versus a randomly chosen subject’s SC matrix (Fig. S9E), there was a substantial improvement in fit for models using a subject’s own optimal synaptic parameters relative to models using randomly chosen subjects’ synaptic parameters (ΔCoD = 0.56; Fig. 2D). Similarly, individual-level synaptic parameters improve fit values over group-level parameters (ΔCoD = 0.26; Fig. 2C), whereas individual-level SC matrices fail to improve fit values relative to group-level SC (ΔCoD= -0.02; Fig. S9F). This suggests that, at least for this modeling framework, the synaptic parameters are predominantly responsible for carrying individual-specific information. We also performed these analyses with the SC processed in three additional ways for tractography seeding and normalization; these processing methods yielded differences in fit values that were generally small (Δ*CoD* ≲ 0.03 on average; Fig. S10), suggesting that the modeling framework is robust to SC processing methods.

### Model captures long-term, subject-specific components of FC matrices

Functional connectivity encapsulates information from a wide range of temporal scales. Some components of FC are temporary and transient while others are stable and subject-specific over long timescales and varying conditions (Finn et al., 2015; Amico and Goñi, 2018; Laumann et al., 2015; Gordon et al., 2017; Miranda-Dominguez et al., 2014; Demeter et al., 2020; Gratton et al., 2018). In the HCP dataset, each subject was scanned four times, with scans 1 and 2 occurring on the same day at approximately the midpoint of the ∼2-hour scanning period and scans 3 and 4 occurring similarly on a different day. To explore the model’s capacity to capture within-subject variation, we fit the model each of the subjects’ scans separately (Fig. 7A). The scan-level simulated FC matrices were each constrained by the group-average SC matrix and fit with scan-level synaptic parameters (Fig. 7A).

**Figure 7.**
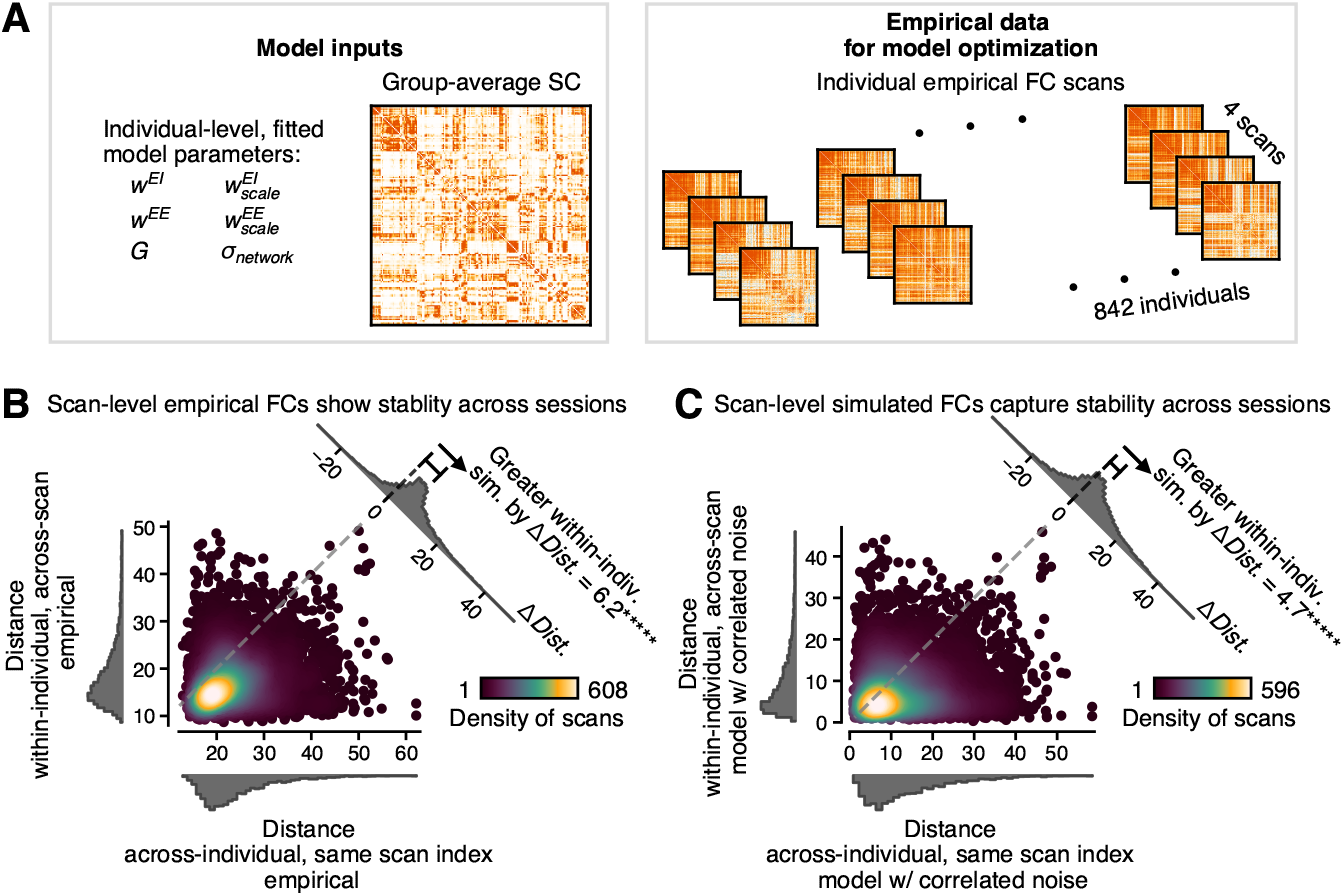
Simulated FC matrices capture long-term, subject-specific components of functional connectivity. **(A)** Schematic showing the empirical inputs to the model. There are 842 subjects with four 15-minute scans, which were modeled using the group-averaged SC matrix. **(B)** Empirical FC matrices are more similar within subject than across subjects. Each scan-level FC matrix was compared to another scan-level FC matrix, either across individuals or across scans. The within-subject, across-scan FC matrix pairs were much more similar than the across-subject, same-scan-index pairs. The similarity metric used here is the the Euclidean distance, meaning more similar matrices have lower values (*****, *p <* 10^−5^, bootstrapping across individuals). **(C)** Same as **(B)**, except the scan-level FC matrices being used are those generated by the 6-parameter heterogeneous model with correlated noise added. While the effect size is smaller, we again found that the within-subject, across-scan FC matrix pairs were significantly more similar than the across-subject, same-scan-index pairs.

For an empirical characterization of within-subject functional variation, we calculated the similarity between pairs of scan-level FC matrices within and across individuals (Fig. 7B and S11A). The similarity function that has been used until now to quantify the distance between predicted and observed values — prediction *r*^2^ or CoD — is asymmetric and not suitable for measuring similarity between two observed values. Thus, we use Euclidean distance as the similarity metric. With Euclidean distance, smaller magnitude values indicate closer similarity; thus, in Fig. 7B the dense collection of points below the diagonal unity line shows that within-subject FC matrices are more similar than across-individual FC matrices. This reaffirms that empirical FC is subject-specific and stable over time.

We used the scan-level simulated FC matrices to test if the model is capable of capturing aspects of empirical within-subject variation. We found a significant tendency for the within-subject simulated FC matrices to be more similar than those across individuals (Fig. 7C). The simulated Euclidean distance values range from ∼ 0–40, where within-subject FC is more similar than across-subject FC (ΔDist. = 4.7, on average). Same-day simulated FC is more similar than different-day FC for individuals (bΔDist. = 1.5, on average) (Fig. S11B,C). This indicates that the model — in particular, the synaptic parameters — is capturing between-subject differences that are not merely due to short-timescale variation in FC.

## Discussion

We used a biophysically-based neural circuit model to simulate resting-state fMRI functional connectivity (FC) of young, healthy adults from the HCP dataset at the level of individual subjects. Our parsimonious model, having only up to six free fitting parameters setting synaptic strengths, was able to capture key aspects of individual variation in FC. Our analyses revealed neurophysiologically interpretable model mechanisms that relate to the three dominant modes of individual variation. We further investigated model fitting of within-subject variation, and we examined potential utility of individual-level structural connectivity (SC). All together, these findings suggest that cortical synaptic dynamics account for a portion of functional variation across subjects, whereas anatomical connectivity – as it can currently be estimated – is insufficient at capturing functional variation across individuals.

Our model captures many features of within- and across-subject variation. Empirically, functional connectivity exhibits subject-level specificity and within-subject stability over time (Finn et al., 2015; Laumann et al., 2015; Gordon et al., 2017; Amico and Goñi, 2018; Miranda-Dominguez et al., 2014; Demeter et al., 2020; Gratton et al., 2018), which are features exhibited by the fitted models both across subjects and across scans within subjects (Figs. 2 and 7). Empirical studies have also indicated that across-subject variation tends to be greater in higher-order association areas than in sensorimotor areas, both anatomically (Hill et al., 2010; Sun et al., 2022) and functionally (Finn et al., 2015; Laumann et al., 2015; Miranda-Dominguez et al., 2014; Demeter et al., 2020; Gratton et al., 2018; Mueller et al., 2013; Xu et al., 2019; Sun et al., 2022; Kong et al., 2018). This is consistent with our findings that the second dominant mode (PC2) of FC variation across subjects is strongly correlated to sensory-association hierarchy (Fig. 3) which is captured by parameterization of hierarchical heterogeneity that varies across subjects.

Functional connectivity is inherently linked with anatomical structural connectivity as the scaffold for long-range interactions across brain regions. FC and SC are correlated at the individual level (Honey et al., 2009), and show some level of subject specificity (Zimmermann et al., 2018a; Messé, 2020). However, in our model individual-level SC did not improve fitting to FC. This finding is consistent with empirical studies seeking to relate individual differences in FC to individual differences in SC, in healthy populations. These empirical studies typically found such SC-FC correlations to be weak and inconsistent (Zimmermann et al., 2018a; Messé, 2020). This limited utility of SC in predicting FC may be due to technical limitations in diffusion tractography at the individual level. Probabilistic diffusion tractography is well known for false positive connections and sensitivity to other biasing factors (Grisot et al., 2021; Zimmermann et al., 2018a; Messé, 2020; Maier-Hein et al., 2017; Jones et al., 2013; Jeurissen et al., 2017; Sotiropoulos and Zalesky, 2019; Zalesky et al., 2016). Of the methods we examined, none meaningfully improved the fit values of the simulated FC at either the group or individual level. Further research is needed to establish a gold standard for non-invasive white matter estimation (Seider et al., 2022; Schilling et al., 2019), which could improve neural circuit models linking SC to FC.

A few studies have previously applied dynamical neural circuit models to BOLD functional connectivity at the individual level. Aerts et al. (2018) simulated FC of 36 individuals, approximately half of whom had been diagnosed with brain tumors, finding that model parameters varied with disease state and behavioral measures. Zimmermann et al. (2018b) modeled the FC matrices of 124 individuals along the spectrum of healthy aging including mild cognitive impairment and Alzheimer’s Disease, finding relationships between model parameters and cognition/disease state. However, to our knowledge, the present study is the first to demonstrate a biophysical model capturing individual differences in FC across a large population of young, healthy individuals. Our study also has a much larger sample size than previously studied, which allowed us to relate model parameterization to the dominant modes of individual variation in FC. A commonality across all of the aforementioned models is a free parameter related to global coupling strength (Zimmermann et al., 2018b; Aerts et al., 2018). This is consistent with our finding that the leading component of variation (PC1) across subjects is strongly correlated with global signal (Fig. 3). However, the previous studies utilized a homogeneous model, thus assuming local synaptic parameters are uniform across the cortex. The present study allows hierarchical heterogeneity across the cortex, which is consistent with the second leading component of variation (PC2) across individuals. This likely explains our model’s ability to effectively capture individual variation across healthy subjects.

Another recent study applied a circuit modeling framework to individual healthy subjects via BOLD time-series data (as opposed to FC matrices, as was done here). developed a data-driven model with a large number of free parameters corresponding to coupled neural masses. This model notably generates subject-specific simulated FC matrices. Compared to our model, this model produces simulated FC matrices that are quantitatively better fit to the empirical data; however, this model is considerably more complex, with over 176,000 free parameters, compared to the 5–6 free parameters used in our model. This allows greater model flexibility, but likely hinders the interpretability and validity of the optimized parameter values relative to a more constrained model (Frässle et al., 2021). Interestingly, the more generalized approach of Singh et al. (2020) finds that optimal parameter values across the cortex follow a hierarchy from sensory to association regions, supporting our characterization of a hierarchical gradient of regional specialization.

There are a number of directions that this model could be extended in the future. For instance, personalized fitting of parameters related to the hemodymanic response function may prove useful, as hemodynamic response functions have been shown to vary across individuals more than across regions (Handwerker et al., 2004). Personalized parcellations and network maps may be especially fruitful to test in future studies. Group-level parcellations can obscure small but meaningful variations across subjects (Gordon et al., 2017), and subject-specific parcellations and network maps are predictive of behavioral measures (Kong et al., 2018). Incorporating additional brain structures, such as thalamic nuclei, into the model may also improve the personalization of the simulations, particularly across disease states. For example, disturbances in thalamo-cortical connectivity are observed in patients with schizophrenia and indicative of individual differences in disease severity (Anticevic et al., 2014).Furthermore, additional brain structures could clarify the origins of model components such as the correlated background noise which could reflect common subcortical inputs. Effects of neuromodulators would also define a promising direction to pursue, which could relate to individual differences and within-subject variability in arousal Pfeffer et al. (2021); Munn et al. (2021).

Beyond our goal of fitting across-subject differences, our approach of expanding the model parametrization based on leading modes of variation could be extended to better study within-subject variation, by leveraging datasets with many scans per subject (Gordon et al., 2017; Noble et al., 2017). Within-subject variability has been shown to follow a different gradient across the cortex and is smaller in magnitude than across-subject variation (Laumann et al., 2015; Gratton et al., 2018). Thus, further investigating the leading modes of within-subject variation of FC across time, brain states, and and task conditions, may provide insight into useful model extensions to better capture the diverse forms of functional variation in the human brain.

## Funding

This study was supported by grants from the National Institute of Mental Health (R01MH112746 and P50MH109429 to JDM; R01MH108590 to AA) and the Swartz Foundation (MD).

## Competing interests

AA and JDM are co-founders of Manifest Technologies, Inc. JLJ is an employee of Manifest Technologies.

## Methods

### Dataset

All data for this study come from the 1200-subject release of the Human Connectome Project (HCP) (Essen et al., 2013). We included all subjects having four scans of resting-state fMRI data, excluding subjects flagged for having anatomical anomalies, for a total of 842 subjects. Subjects were aligned to a common atlas using the Multimodal Surface Matching (MSMAll) algorithm (Robinson et al., 2014). The cortex was partitioned into 360 parcels (180 per hemisphere) according to the multi-modal parcellation developed by Glasser et al. (Glasser et al., 2016). For some analyses, the parcels were further grouped into functionally-defined networks (Ji et al., 2019).

#### 6.1.1 Resting-state functional connectivity processing

Resting-state fMRI time series data for each subject were processed according to the HCP minimal processing pipeline (Glasser et al., 2013) and further cleaned using ICA-FIX (Griffanti et al., 2014) (Salimi-Khorshidi et al., 2014), which uses independent component analysis to separate signal from noise. Motion scrubbing was performed to remove frames flagged for badness, consisting of any frames having a framewise displacement greater than 0.5 *mm* or median normalized image intensity root mean squared error greater than 1.6 (Power et al., 2012). The time series data were parcellated, and functional connectivity (FC) matrices were calculated as the Pearson correlation coefficient between time series data for pairs of parcels.

#### 6.1.2. Structural connectivity processing

We obtained minimally pre-processed diffusion-weighted MRI data for each of 328 unrelated (non-twin and non-sibling) adults from the HCP dataset (Essen et al., 2013; Sotiropoulos et al., 2013). FSL’s Bedpostx was then used to estimate the diffusion parameters of up to three crossing fiber orientations for each voxel using a model-based deconvolution approach with zeppelins (Jbabdi et al., 2012; Sotiropoulos et al., 2016), using the following parameters: burn-in period of 3000, 1250 jumps (sampled every 25), automatic relevance determination, and rician noise. Whole-brain probabilistic tractography was then obtained from FSL’s probtrackx pipeline (Behrens et al., 2007; Hernandez-Fernandez et al., 2019), which generated a dense (voxel-by-voxel), distance-corrected structural connectivity matrix. This pipeline seeds from each grey-ordinate 10,000 times, initiated to sample a sphere around the center of the grey-ordinate, to reconstruct unidirectional grey-ordinate to grey-ordinate connectivity. The processing software refers to this seeding strategy as Matrix1 or Conn1. Successful streamlines stopped when they entered the pial surface or entered and exited subcortical grey matter. Tractography was constrained with a fiber threshold of 0.01 and a curvature threshold of 0.2 (which corresponds to minimum angle of approximately ±80 degrees). Streamlines were also discarded if they looped back onto themselves or traveled more than 2000 steps (equivalent to 1 meter with a step length of .5mm). This results in a 91,282-by-91,282 cortical and subcortical structural connectivity matrix. The matrix was then symmetrized by averaging it with its transpose.

The dense structural connectivity matrices were waytotal-normalized by dividing the streamline counts for each grey-ordinate by the waytotal value (i.e., the total number of valid streamlines for each subject) at the individual level to account for streamline magnitude differences across subjects. The dense 91,282-by-91,282 structural connectivity matrices were then parcellated with a multimodally defined cortical parcellation (Glasser et al., 2016), to generate individual-level cortical structural connectivity matrices of size 360-by-360. The structural connectivity matrices were also averaged across subjects to create a group-average structural connectivity matrix. As part of the modeling procedure, all structural connectivity matrices were row-sum normalized such that each matrix row sums to 1, except where noted (Fig. S10C,F).

The structural connectivity matrices were also processed in a few alternate ways to test whether similar results are achieved for different processing strategies (Fig. S10). One variant used the “Conn3” seeding strategy, where FSL’s probtrackx probabilistic tractography pipeline seeds from each white matter voxel to assess bidirectional voxel to grey-ordinate connectivity, which is then used to reconstruct grey-ordinate to grey-ordinate connectivity. This seeding strategy is referred to in the processing software as Matrix3 or Conn3. For another variant, the matrices of streamline counts were log-transformed before being row-sum normalized. Before log-transforming, the individual-level streamline count matrices were normalized to sum to 1*e*18 to account for variations in the number of total valid streamlines across subjects. For a third variant, the structural connectivity matrices were row-and-column-sum normalized such that each matrix sums to 360. For each variant, all other processing steps were carried out as previously described.

#### 6.1.3. T1w/T2w map processing

Maps of the ratio of T1-to T2-weighted (T1w/T2w) images were bias field-corrected and parcellated. Each parcel in the left hemisphere was paired with an analogous parcel in the right hemisphere, and the T1w/T2w maps were symmetrized using the average values between the pairs of parcels.

### Circuit Model Scheme

This study employs a low-dimensional cortical circuit model developed in part by Demirtaş et al. whose free parameters represent circuit-level activity, allowing the macroscopic across-individual variations apparent in fMRI scans to be understood in terms of the underlying cellular architecture (Demirtaş et al., 2019) (Wong and Wang, 2006) (Deco et al., 2014). This model treats the cortex as a network of interconnected nodes, representing each cortical parcel as a node. Each parcel is approximated as a circuit consisting of one excitatory unit (representing pyramidal neurons) and one inhibitory unit (representing interneurons) (Fig. 1A). The excitatory units interact via long-range interactions with each other as well as locally with their respective inhibitory unit and recurrently. The inhibitory units interact locally with their respective excitatory unit and recurrently. This system can then be described by a set of differential equations, which can be solved to reveal a matrix of correlations between the parcels, i.e., a simulated FC matrix (Fig. 1B).

In particular, this system is described by the following set of nonlinear differential equations (Deco et al., 2014):

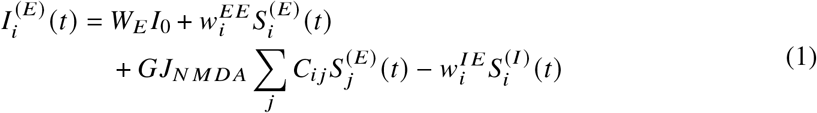

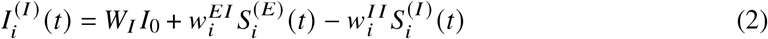

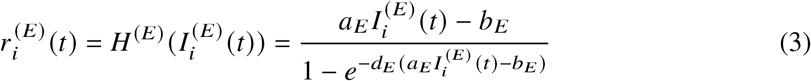

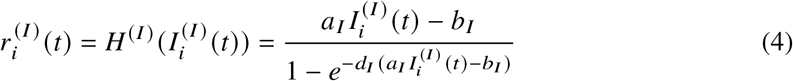

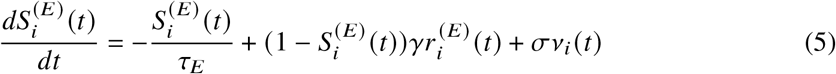

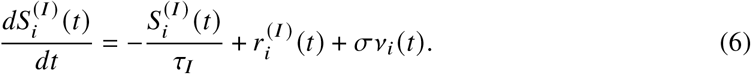

The sub- and superscripts *E* and *I* signify excitatory and inhibitory populations, respectively. 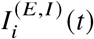 are the input currents to the excitatory/inhibitory units in cortical parcel *i. I*_0_ is the total effective external input current and is set to 0.382*n A. W*_*E,I*_ are scaling factors for *I*_0_ for the excitatory and inhibitory units, respectively. 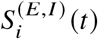 denote the average synaptic gating variables of parcel *i*, representing the fractions of open neuronal channels. *J*_*NMDA*_ is the excitatory synaptic coupling and is set to a value of 0.15*n A. C* represents the structural connectivity (SC) matrix such that *C* is the anatomical connectivity between parcels *i* and *j*. The firing rate of parcel *i* is given by 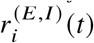. The functions *H* ^(*E,I*)^ () represent neuronal input-output functions, which convert input currents into neuronal firing rates. The parameters *a*_*E,I*_, *b*_*E,I*_, and *d*_*E,I*_ are gating variables and are set to constant values. The parameters *τ*_*E,I*_ represent the decay constants for NMDA and GABA synapses such that *τ*_*E*_ = *τ*_*N MDA*_ = 0.1*s* and *τ*_*I*_ = *τ*_*ABA*_ = 0.01*s*. The kinetic conversion factor *γ* is set to a value of 0.641. Input noise in the system is represented by *v*_*i*_ *(t)*, which in the simplest case is assumed to be a spatially uncorrelated Gaussian process, and is scaled by *σ* = 0.01*n A*. Full model description is presented in Demirtaş et al. (2019).

The parameters 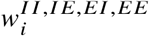 and *G* represent synaptic strengths and embody the variation across individuals in this model. The parameters 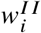 are arbitrarily set to a value of 1.0 for all parcels, and 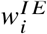 are set to maintain a uniform baseline firing rate of ∼ 3 Hz (with the exact value depending on other parameters of the model). The parameters 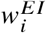 and 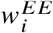 represent the strengths of the local E-I and E-E interactions within parcel *i*, respectively. The global coupling parameter *G* represents the strength of long-distance across-parcel connections, acting as a scaling factor on the SC matrix. Assuming all parcels across the cortex have homogeneous synaptic strength values such that 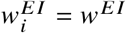 and 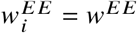 for all parcels defines the homogeneous circuit model. The three free parameters in this model are then *w*^*EI*^, *w*^*EE*^and *G*.

Demirtaş et al. extended the former model to a heterogeneous circuit model, allowing each parcel *i* across the cortex to have different values of 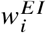 and 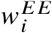. In particular, heterogeneity was incorporated by assuming 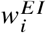 and 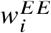 vary linearly across the cortex according to hierarchy, using linearized T1-weighted/T2-weighted (T1w/T2w) maps as a proxy for cortical hierarchy (Burt et al., 2018) (Fig. 1A). Thus, 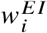 and 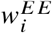 are simplified to being described by two free parameters each: *w*^*EI*^ and *w*^*EE*^ which give the local synaptic strengths in the parcel at the lowest end of the cortical hierarchy, and 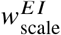 and 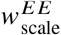, which scale *w*^*EI*^ and *w*^*EE*^, respectively. Thus, local synaptic strengths are linearly increased for parcels higher on the cortical hierarchy. The five free parameters in this model are then 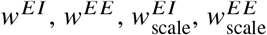, and *G*. Demirtaş et al. found that this heightened flexibility improved the model’s ability to reproduce empirical measures of dissimilarity across subjects (Demirtaş et al., 2019), suggesting that allowing the model parameters to exhibit heterogeneity across the cortex may be especially important for modeling at the individual subject level.

These differential equations are further constrained using the Balloon-Windkessel model of hemodynamic response to generate a simulated BOLD signal from the synaptic gating variables 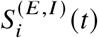 (Friston et al., 2003) (Deco et al., 2013). Defining **P** as the covariance of synaptic gating variables,

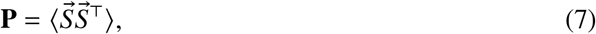

we obtained the relation:

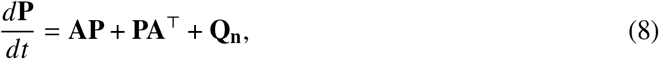

where **A** is the Jacobian matrix and **Q**_**n**_ is the noise covariance matrix. This may be evaluated at a stationary state, yielding:

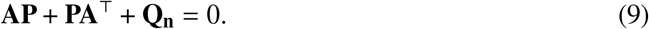

This expression is a Lyapunov equation and is readily solvable, thus generating a simulated FC matrix. Further details are provided in Demirtaş et al. (2019). While this model is equally applicable to whole brain FC matrices, we chose to model only the left hemisphere of the cortex to lessen computational load.

#### 6.2.1 Network-structured correlated noise model extension

Noise is a time-varying quantity represented by *v*_*i*_ (*t*) and scaled by *σ* in Eqs. 5 and 6. In the simplest case, noise input to the synaptic gating variables is spatially uncorrelated, making the noise covariance matrix **Q**_**n**_ identical to the identity matrix (Figs. 2 & 4). We proposed an extension to the model where same-network parcels have correlated input noise between their E-E connections.

Thus, we added a correlated noise matrix to the identity **Q**_**n**_ matrix, which has a block structure for same-network, E-E connections. Here we used functional network assignment from Ji et al. (2019). In the same way that the spatially uncorrelated noise is scaled by *lT*, the within-network correlated noise is scaled by a sixth free parameter *lT*_network_, which is optimized in the same way as the other free parameters.

### Parameter optimization algorithm

The free parameters — *w*^*EI*^, *w*^*EE*^ and *G* for the homogeneous model, 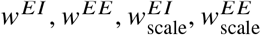, and *G* for the heterogeneous model, or 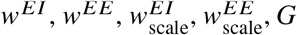, and *σ*_network_ for the correlated noise heterogeneous model — are fitted at the single-subject level. In particular, a differential evolution algorithm was used to search for optimal parameter values via iteratively choosing sets of parameters semi-stochastically within a given domain of possible parameter values. This algorithm is useful for finding the global extrema of multidimensional functions that may be discontinuous or noisy. Using these randomly-generated parameter values, our modeling algorithm calculates a simulated FC matrix and computes its similarity to the respective empirical FC matrix. After a large number of iterations, it eventually converges to the set of parameters maximizing the similarity between the simulated and empirical FC matrices. Periodically, the differential evolution algorithm found multiple solutions which fit the empirical data similarly well. In these cases, the model was run multiple times, and we chose the simulated FC (and associated parameter values) maximally fitting the empirical FC.

The parcellation used here contains 180 parcels per hemisphere. Thus, the simulated FC matrices (which contain only the left hemisphere) are of size 180 × 180. These matrices are symmetrical and have a value of 1 for all entries along the diagonal (as the time series data for a given parcel is perfectly correlated with itself). Thus, each 180 × 180 simulated FC matrix contains 16,110 non-trivial, unique values. By contrast, our models incorporate between three and six free parameters. Based on the fact that the FC-space used here has a dimensionality ∼ 2,000 times that of the free parameter space, we posited that overfitting is unlikely to negatively impact fit validity. We tested this with an exemplification of the models used in this study: the 5-parameter heterogeneous model simulating the FC matrices of individual subjects. We split up each individual’s time series data into training and testing data in two ways: by assigning alternating time points to the training/testing sets and by splitting each of the four scans into quarters and assigning alternating quarters to the training/testing sets. These training and testing time series data were then used to create training and testing FC matrices (as described previously). The optimal modeling parameters for each subject were found by simulating the training FC matrices. The fit values were then calculated between the subject-level simulated training FC matrices and each of the empirical training and testing FC matrices (Figs. S1F,G). Splitting via every other time point, the fit values between training and testing data are virtually indistinguishable, with the training data producing fit values that are higher and only weakly significant (Fig. S1F). Splitting the scans into quarters, the training data is fit significantly better than the testing data; however, the vast majority of subjects’ training and testing data are fit similarly well such that the effect size between training and testing is considerably less than effect sizes seen in previous results (Fig. S1G). The outliers for whom the difference between training data and testing data are large could be suggesting that the model is capturing short timescale phenomena, which are physically meaningful and could be further studied. Overall, overfitting does not seem to be causing artificially high fit vales, so we opted not to hold out any data.

### Choice of similarity metric

Previous studies have primarily used the Pearson correlation coefficient to quantify the similarity between simulated and empirical data (Zimmermann et al., 2018b; Aerts et al., 2018, 2020; Demirtaş et al., 2019; Wang et al., 2019; Deco et al., 2013). However, this metric overestimates significance for weakly correlated variables (Scheinost et al., 2019), which is problematic, as we are operating in the regime of relatively low correlation values. Moreover, the Pearson correlation coefficient is blind to differences in slope or intercept, potentially finding substantially dissimilar FC matrices to be highly correlated (Fig. S1A-C).

Thus, as suggested by Scheinost et al. (2019), we chose to primarily perform our fitting procedures and subsequent analysis using the measure “prediction *r*^2^” defined as:

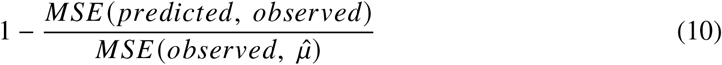

where 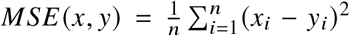, *predicted* refers to the simulated FC matrix, *observed* refers to the empirical FC matrix, and 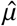 is the average value of the empirical FC matrix. Prediction *r*^2^ values fall into the range, [−∞1], with a value of 1 implying a perfect correlation, a value of 0 implying the model fits as well as a constant model whose value is the mean of the data, and negative values implying arbitrarily poorly fitting models. For highly correlated sets of data, prediction *r*^2^ values converge to the square of the Pearson correlation coefficient (Fig. S1D,E). To maintain a clear distinction between the various definitions of *r*^2^ and to relieve potential confusion due to displaying negative *r*^2^ values, we chose to use the generic term “coefficient of determination” (CoD) to refer to the metric used for fitting, which in the present study is synonymous with “prediction *r*^2^”.

### Example individuals

In some instances, we explicitly compared the FC matrices of specific individuals (Fig. 2). To that end, we selected two individuals who were well-fit by the individual-level model, one of whom was fit similarly well by the group-level model (individual A) and one of whom was fit poorly by the group-level model (individual B). The HCP subject IDs given to these subjects correspond to 833148 for individual A and 112920 for individual B.

### Principal components of variation within and across individuals

Principal component analysis (PCA) provides a window to study variation across and within individuals. For PCA across subjects, we defined FC space as the feature space and the 842 individuals as the samples. Since the FC matrices are symmetric, we flattened the upper triangle of the individual-level FC matrices to generate a vector of “features” in FC space for each subject. We performed PCA across individuals, yielding vectors in FC space representing the modes of FC variation across subjects. We then reshaped these vectors into square, symmetric matrices (of shape 180 ×180) for consistency with the FC matrices. For within-subject PCA, we again defined FC space as the feature space. The samples are defined as tenths of a scan. To that end, we divided each individual’s BOLD timeseries data into ten segments and generated the FC matrix for each segment. This yields 40 FC matrices for each subject (since each subject had four scans), each of which is based on /*sim*1.5 minutes of data. For each individual, we again flattened the upper triangle of each FC matrix and performed PCA across tenths of a scan. This yields the modes of within-subject FC variation. As before, these are then reshaped into symmetric matrices of shape 180 × 180 for each subject.

#### 6.6.1 Calculation of statistical significance

Figure 3H and Supplementary Figure S4A depict the similarities between the leading PCs of across-subject variation and various topographical features. Statistical significance was tested by bootstrapping across subjects to sample the leading PCs a large number of times. From this, we estimated distributions of correlations between PCs and topographical features and calculated significance under the null hypothesis that a given PC is equally correlated with two given neural features.

### Alignment of PCs in sensitivity subspace

The modeling algorithm returns the simulated FC matrix that maximally fits the empirical data, as well as the optimal parameter values corresponding to this matrix. These optimal parameter values may be represented as 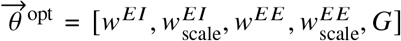 for the heterogeneous model or 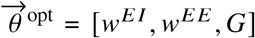 for the homogeneous model (Fig. 4A, left panel). Each subject has individualized optimal parameters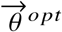for each model (heterogeneous or homogeneous). For each subject, one of the optimal parameters 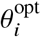 was perturbed away from its optimal value by adding (or subtracting) a small amount to (or from) its value, resulting in 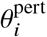. We then simulated the FC matrix corresponding to the set of optimal parameters with 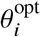 replaced by 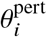, which we denoted by 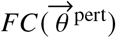. We defined the *θ*_*i*_ sensitivity matrix as the difference between the optimal simulated FC matrix 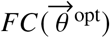 and the perturbed simulated FC matrix 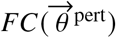 divided by the difference between 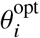 and 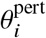. This could also be expressed as 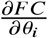 (Fig. 4A, center panel). This is repeated for each of the five (or three) parameters of the heterogeneous (or homogeneous) model (example sensitivity matrices for the heterogeneous model for one subject shown in Fig. S4B).

The upper triangle of the sensitivity matrices can be flattened into vectors, the linear combination of which define a low-dimensional subspace of the much larger dimensionality FC-space. In particular, this sensitivity subspace represents a subject-level estimate of the portion of the ∼16,000-dimensional FC-space that is capable of being produced by the model, estimated local to the set of subject-specific optimal parameters.

We then projected each of the flattened leading empirical PCs (Fig. 3B–D) onto this subspace (Fig. 4A, right panel). This projection is normalized relative to the length of the PC vector, yielding a value between 0 and 1, which we defined as an alignment. Since the leading PCs encode the dimensions across which individual subjects vary, the alignment of the leading PCs within the sensitivity subspace serves to quantify how well each of the modes of across-individual variation can be captured using a given model (heterogeneous or homogeneous, in this case). Thus, an alignment value of 1 implies that the model is able to entirely capture that mode of variation across subjects, while an alignment of 0 implies the model is blind to that mode of variation.

**Figure S1.**
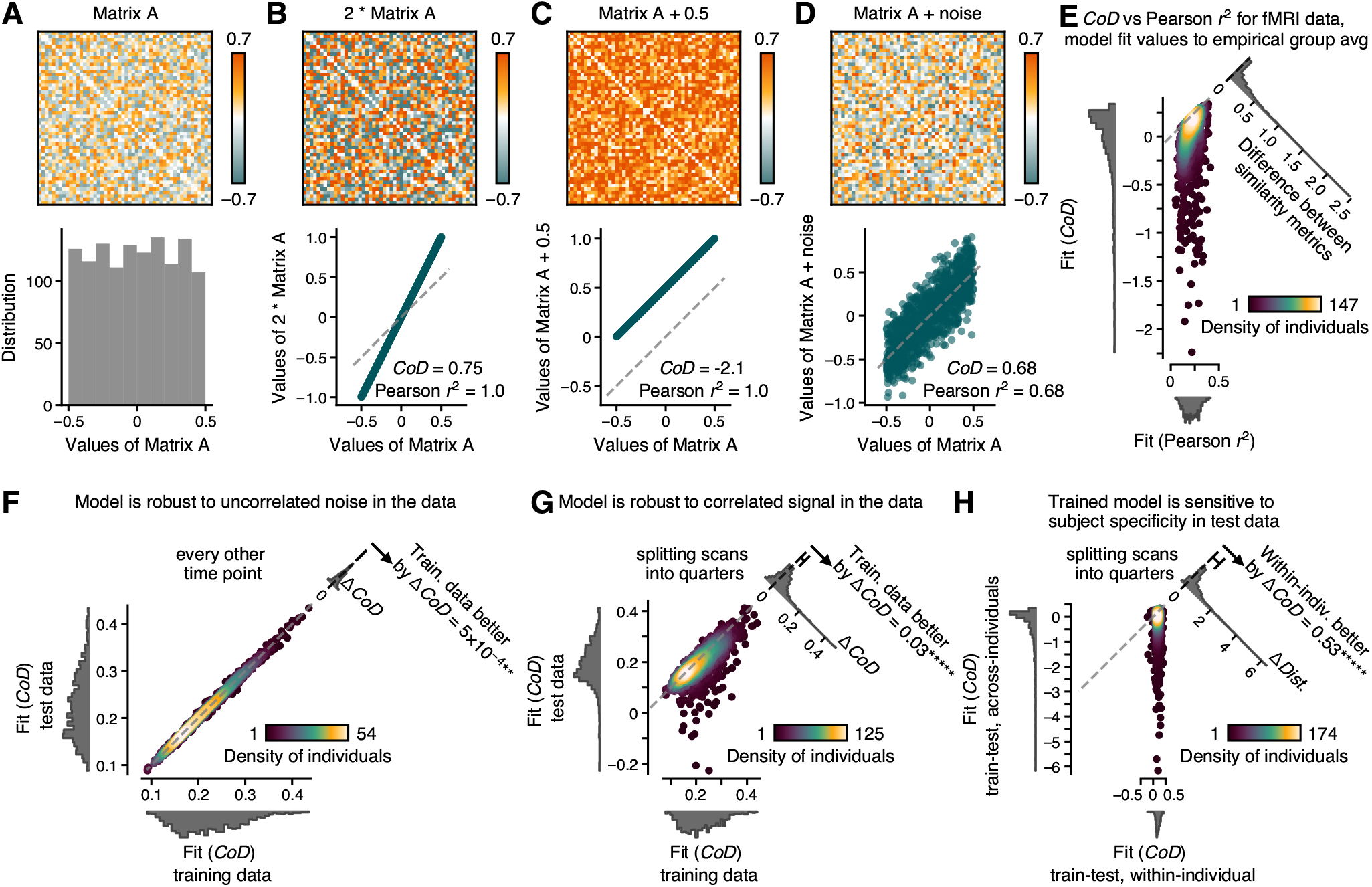
Prediction *r* ^2^ (referred to here as coefficient of determination, CoD) provides more biophysically meaningful estimate of similarity between simulated and empirical FC matrices (Scheinost et al., 2019). **(A)** Randomly generated symmetric matrix of values distributed uniformly between -0.5 and 0.5. All matrices shown are symmetric and have values within [-1, 1], representing possible correlation matrices. **(B)** Pearson correlation is blind to differences in slope, while CoD sees a reduction of similarity. **(C)** Pearson correlation is also blind to differences in intercept, which produce correlation matrices that are physically very distinct. CoD is sensitive to differences in intercept and returns a low fit value accordingly. **(D)** For the case of highly correlated variables, CoD converges to the square of the Pearson correlation coefficient. **(E)** Comparing the correlations between empirical individual-level FC matrices and the simulated group-averaged FC matrix for two different similarity metrics: CoD and the square of the Pearson correlation coefficient. The Pearson correlation coefficient inflates similarity, especially for poorly fit subjects. Well-fit subjects have similar fit values using either metric. **(F)** Evaluating the model’s ability to fit held out test data with simulated FC matrices optimized to training data. Holding out every other time point to generate test/training FC matrices, simulated FC matrices optimized to the training data fit the test data as well as the training data (**, *p* = 0.007, bootstrapping across individuals). **(G)** Splitting the time series data into four equal portions and generating test and training FC matrices using alternating quarters shows a small but significant improvement in the simulated FC matrices’ fits to the training data relative to the test data. This suggests there may be time-correlated signal within the data, to which the model is sensitive (*****, *p <* 10^−5^, bootstrapping across individuals). **(H)** Same as Fig. 2D, except fits are calculated between the trained model and the held-out test data, where test and training data are defined as in panel G. This suggests that the effect seen in Fig. 2D is not due to overfitting (*****, *p <* 10^−5^, bootstrapping across individuals).

**Figure S2.**
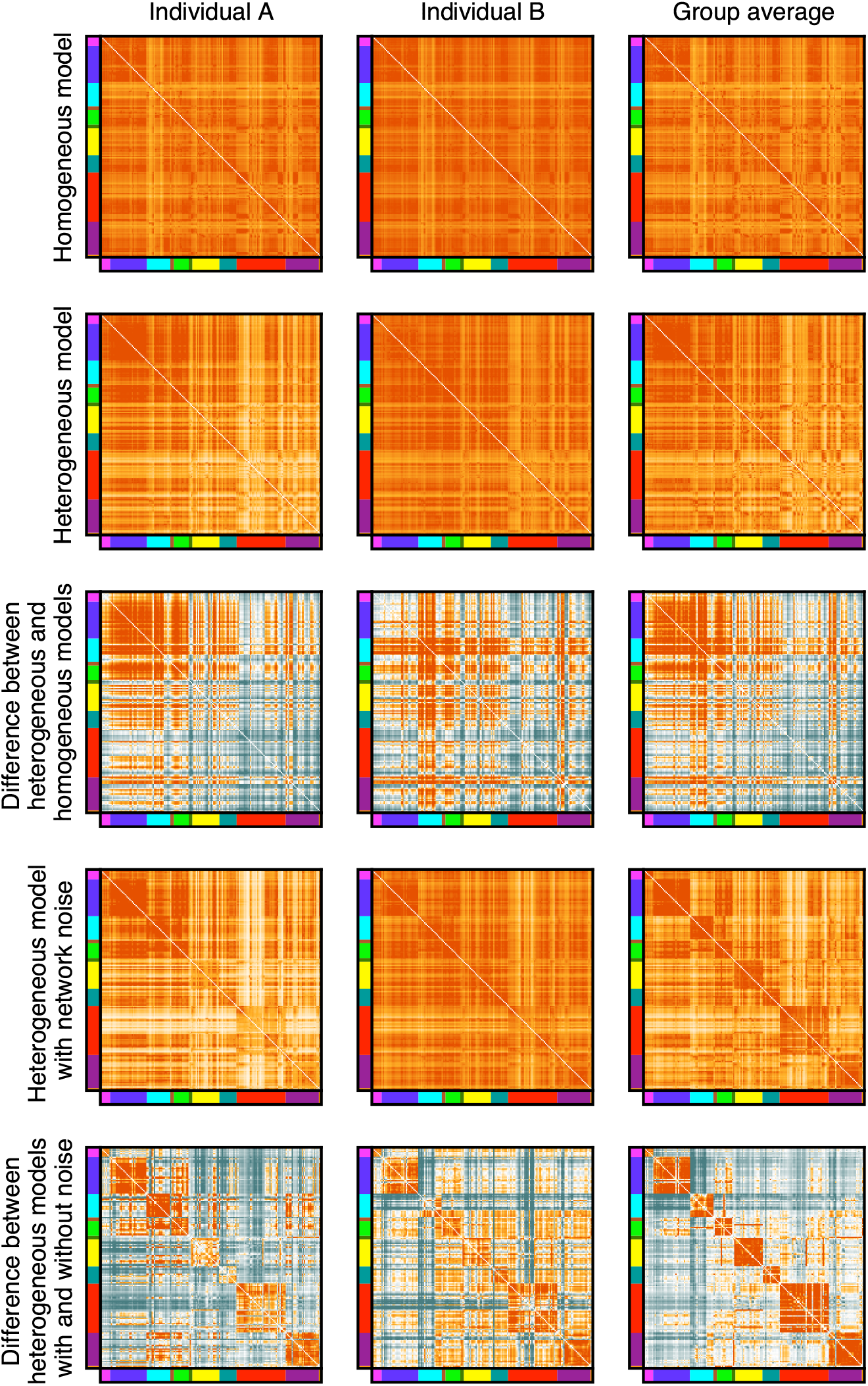
Simulated FC and difference matrices for two subjects. and the group average. The difference between the heterogeneous and homogeneous models shows distinct hierarchical structure. Similarly, the difference between the heterogeneous model with and without within-network noise shows distinct network-like structure.

**Figure S3.**
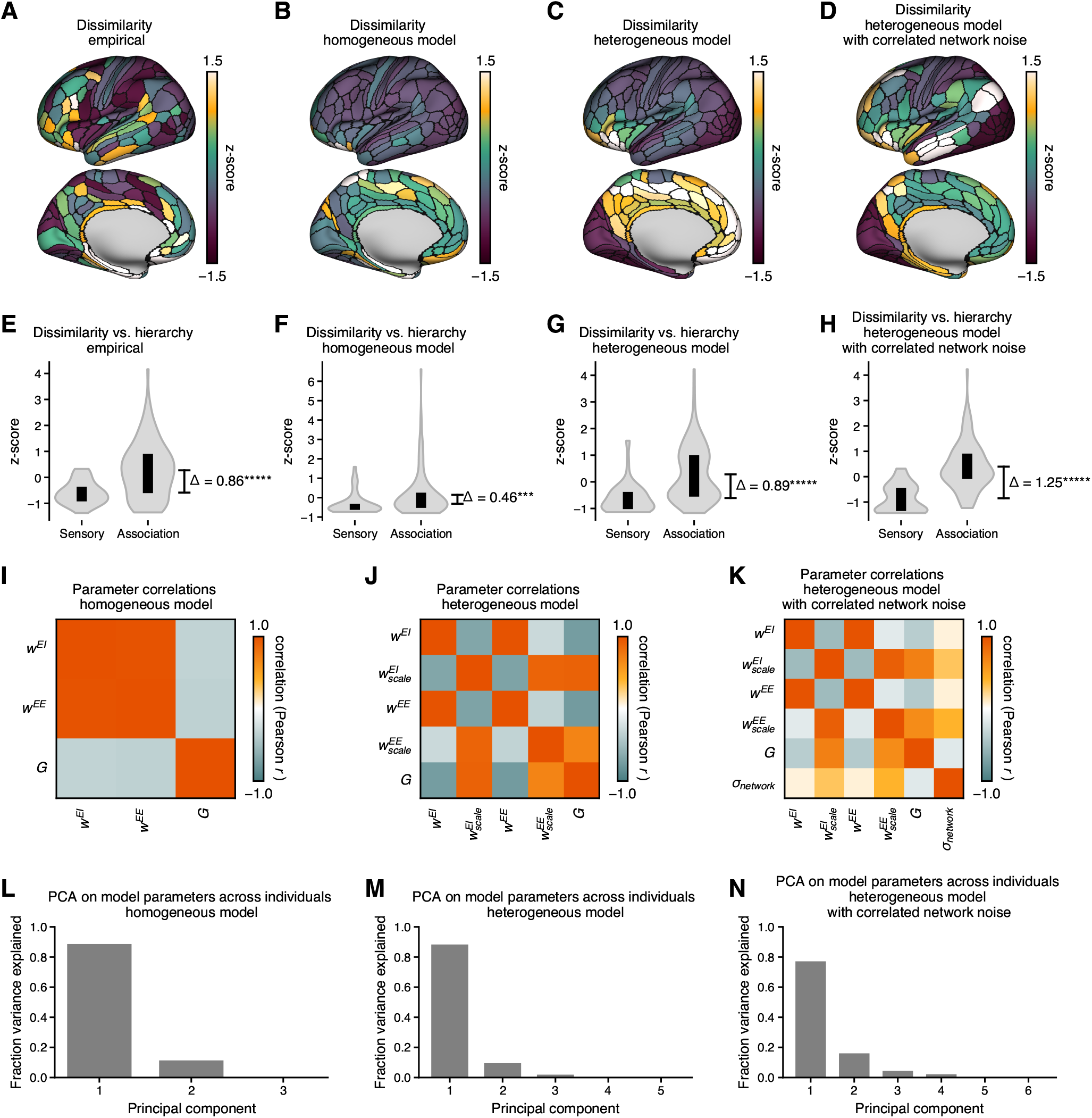
Characterization of empirical and simulated across-individual dissimilarity and parameter correlations. **(A, E)** Topography of dissimilarity across subjects for empirical FC matrices (as defined in Fig. S9), following the example of Mueller et al. (2013). Like the cortical hierarchy map (Fig. 3F), this shows relatively low dissimilarity across sensorimotor regions and high dissimilarity across association regions. (*****, *p <* 10^−5^, bootstrapping across parcels) **(B, F)** Same as Panels A & E, except for the homogeneous model. Some disparity is seen between sensory and association regions; however, the effect is less pronounced than in the empirical data. (***, *p <* 10^−3^, bootstrapping across parcels) **(C, G)** Same as Panels A & E, except for the heterogeneous model. This model shows a disparity between sensory and association that is similar to the empirical data. (*****, *p <* 10^−5^, bootstrapping across parcels) **(D, H)** Same as Panels A & E, except for the heterogeneous model with correlated network noise added. This model also shows a disparity between sensory and association that is similar to the empirical data. (*****, *p <* 10^−5^, bootstrapping across parcels) **(I)** Correlations between the optimal individual-level parameters for the homogeneous model. The weight parameters *w*_*EI*_ and *w*_*EE*_ are highly degenerate with each other, as evidenced by the high correlation value between them. **(J)** Same as Panel I, except for the heterogeneous model. The weight parameters *w* ^*EI*^ and *w* ^*EE*^ are still highly degenerate, while the added scaling parameters 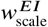 and 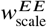 show less degeneracy. **(K)** Same as Panel I, except for the heterogeneous model with correlated network noise added. The additional parameter *σ*_network_ is not strongly correlated with any of the other parameters, justifying the addition of this parameter. **(L)** Analysis of the degeneracy of the parameters. PCA was performed across individuals on the matrix of optimal fit parameters for each subject. The variation in the model parameters is entirely captured by two dimensions, which results from the degeneracy between the parameters *w* ^*EI*^ and *w* ^*EE*^. **(M)** Same as Panel L, except for the heterogeneous model. The variation in the model parameters is captured by three dimensions. **(N)** Same as Panel L, except for the heterogeneous model with correlated noise added. The variation in the model parameters is captured by four dimensions.

**Figure S4.**
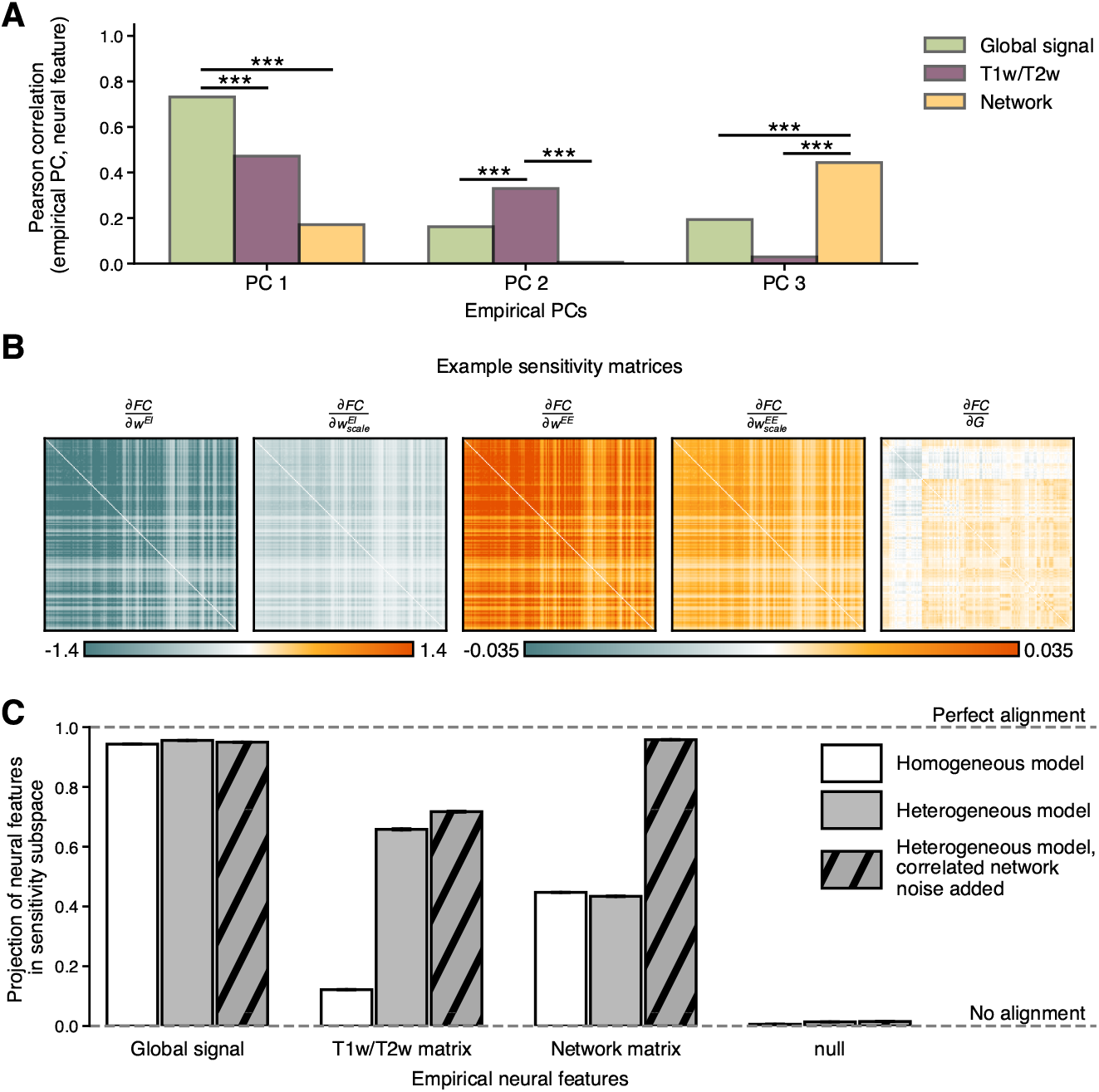
Supplement to Figs. 3, 4, & 5. **(A)** Pearson correlation coefficient between leading three principal components (Figs. 3B–D) and neural features (Figs. 3E– G). Consistent with results found using cosine similarity, PC1 is highly correlated with global signal, significantly more than both T1w/T2w and network structure. PC2 is highly correlated with hierarchy (significantly more than both global signal and network structure), and PC3 is highly correlated with the network structure (significantly more than both global signal and hierarchy) (***, *p <* 10^−3^, calculated by bootstrapping across individuals). **(B)** Example sensitivity matrices for one subject for the heterogeneous model. Within the local neighborhood of the optimal parameters, there is degeneracy apparent across the weight parameters (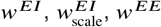, and 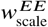). **(C)** The alignment of various neural features (global signal, T1w/T2w matrix, network matrix; shown in Figs. 3E–G) onto the sensitivity subspaces for individual subjects for each of the homogeneous, heterogeneous, and heterogeneous with correlated noise models. Global signal aligns closely with the sensitivity subspaces for all three models. The T1w/T2w matrix aligns much more closely with the subspaces for the two heterogeneous models. The network matrix aligns much more closely with the heterogeneous model with correlated noise than the homogeneous or heterogeneous models. These results are expected, given that heterogeneous models directly incorporate T1w/T2w maps and the correlated noise was generated with a network structure. This serves as a proof of concept that this novel method of projecting FC-like matrices onto the sensitivity subspaces returns meaningful estimates of the model’s ability to capture features of the projected matrix.

**Figure S5.**
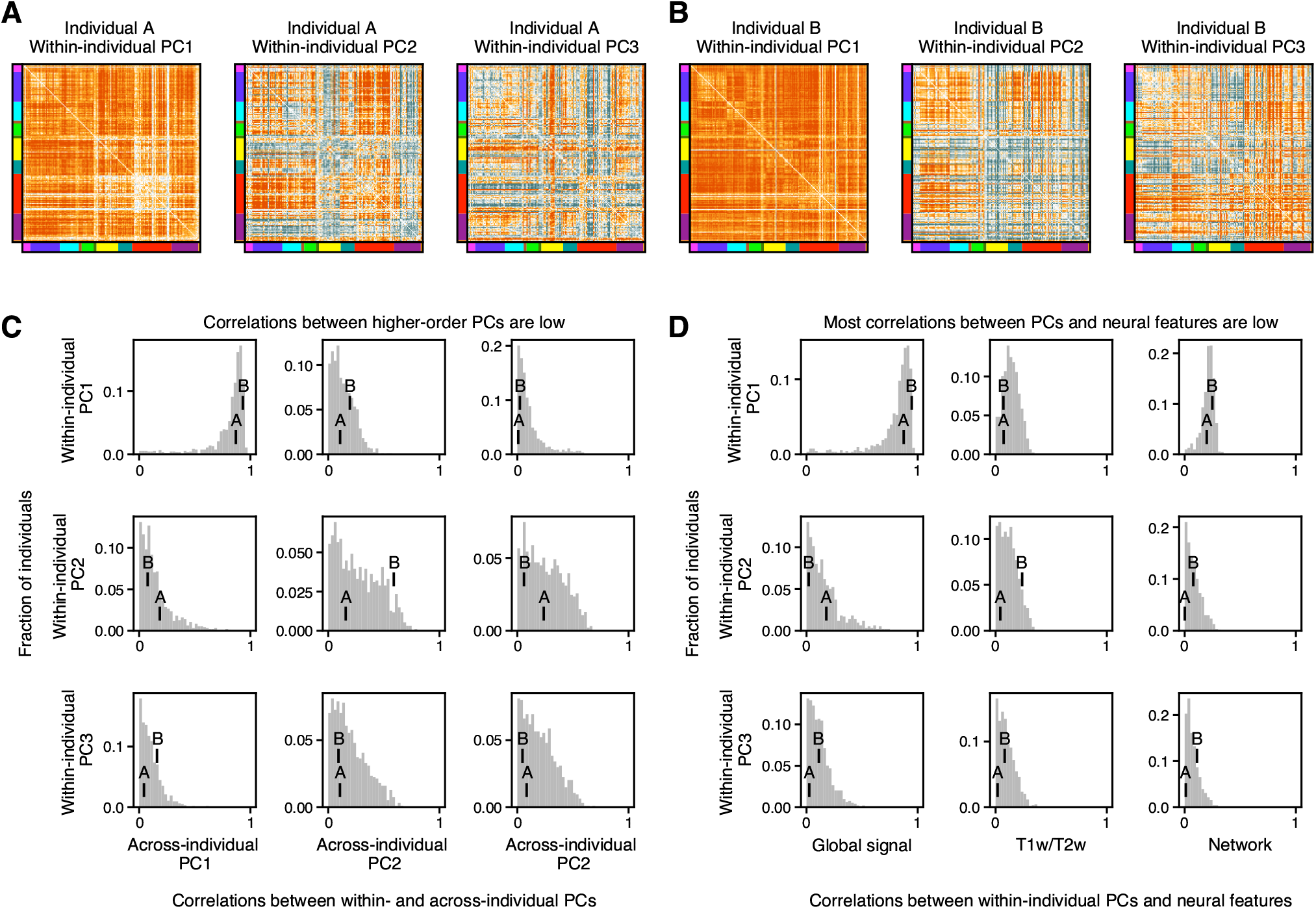
In general, functional within-subject variation empirically behaves differently than variation across individuals. **(A, B)** First three leading principal components of within-subject variation for two example individuals. **(C)** Correlation between within- and across-individual variation for all subjects. For most individuals, within-subject PC1 is highly similar to across-subject PC1. However, correlations between higher-order PCs are generally much weaker. **(D)** Correlation between within-subject variation and previously explored neural features for all subjects. Similar to (C), within-subject PC1 is highly correlated with the global signal matrix for most individuals, while correlations involving the higher-order PCs and other neural features tend to be much weaker.

**Figure S6.**
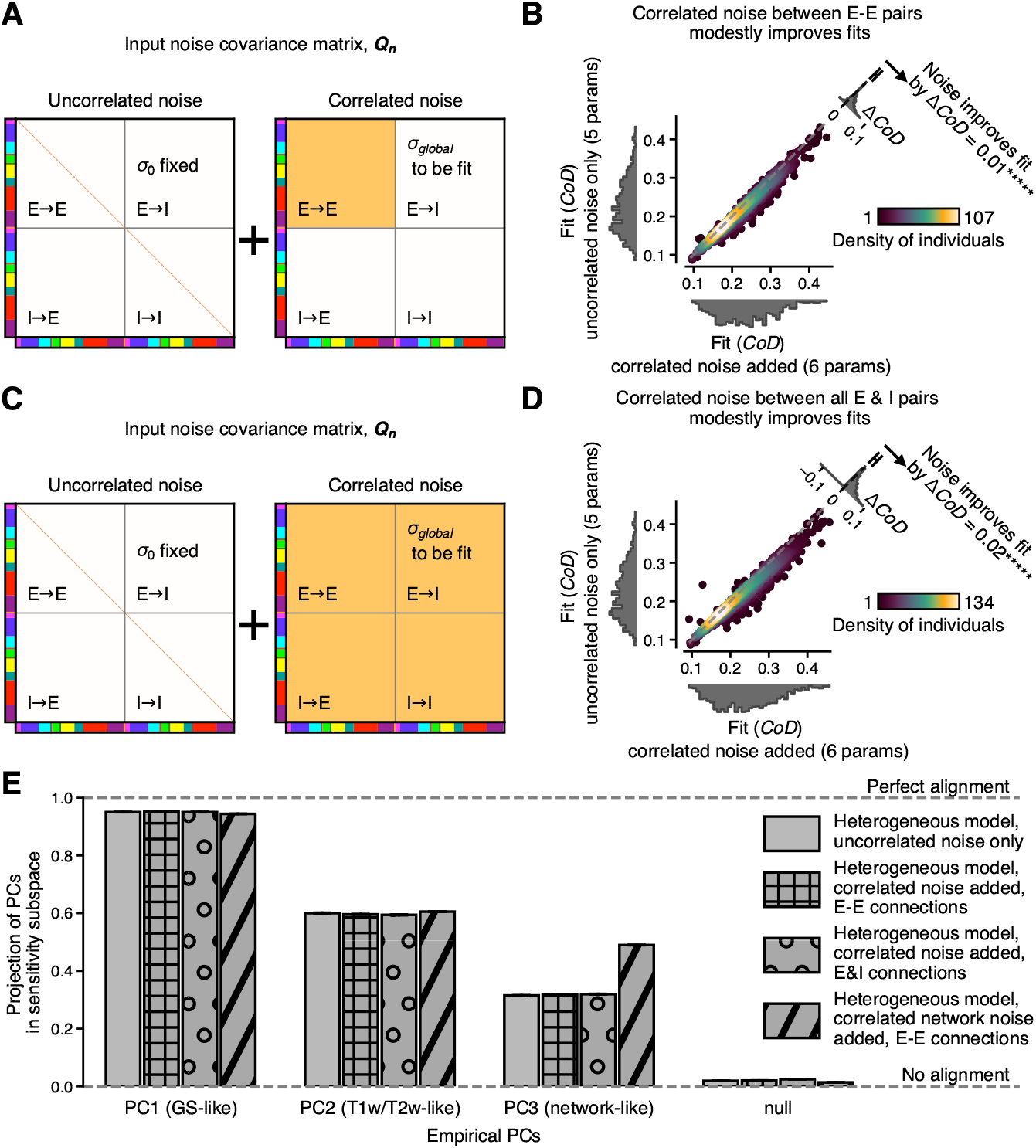
Supplement to Fig. 5 showing alternate extensions to the model. **(A)** Schematic showing addition of correlated noise between all pairs of excitatory units. **(B)** A model with correlated noise between pairs of excitatory units modestly improves fits to empirical data. (*****, *p <* 10^−5^, bootstrapping across individuals) **C)** Schematic showing addition of correlated noise between all excitatory and inhibitory units. **(D)** A model with correlated noise between all units modestly improves fits to empirical data. (*****, *p <* 10^−5^, bootstrapping across individuals) **(E)** Alignment of leading principal components of individual variation onto the subspace of sensitivity matrices for four models: the heterogeneous model with uncorrelated noise, the heterogeneous model with correlated noise (Fig. S6A), the heterogeneous model with correlated noise (Fig. S6C), and the heterogeneous model with correlated noise (Fig. 5A) for comparison. Incorporating correlated noise in these ways modestly improves fit values but fails to capture PC3 as well as with correlated within-network noise (Fig. 5).

**Figure S7.**
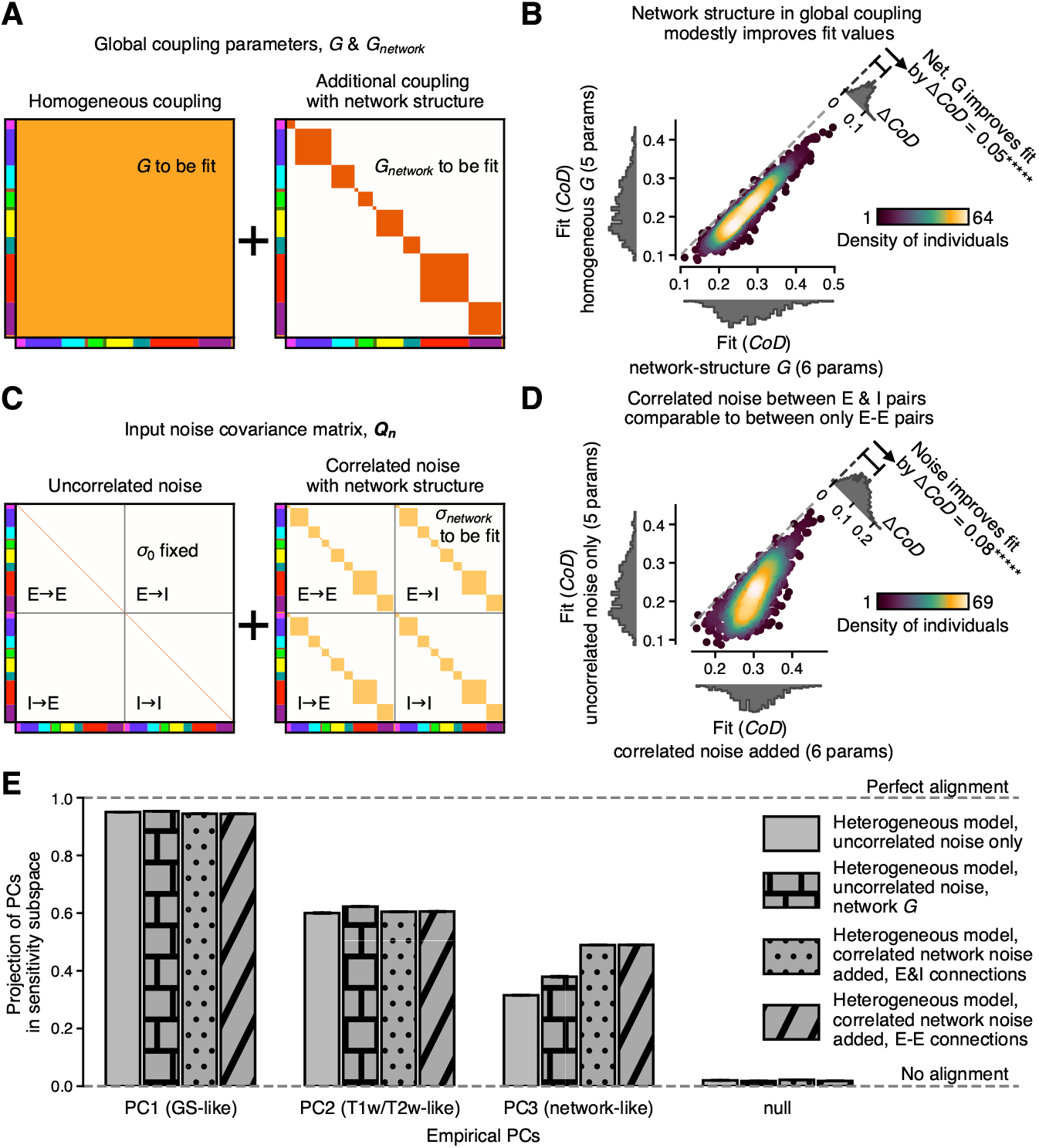
Supplement to Fig. 5 showing alternate extensions to the model incorporating network structure. **(A)** Schematic showing addition of network structure to global coupling parameter *G*, allowing stronger coupling between within-network units. **(B)** A model with network structure in global coupling modestly improves fits to empirical data. (*****, *p <* 10^−5^, bootstrapping across individuals) **(C)** Schematic showing addition of network-structured correlated noise between pairs of excitatory and inhibitory units. **(D)** A model with correlated network noise between E & I units improves fits to empirical data in a similar manner as a model with correlated network noise between only E-E connections. (*****, *p <* 10^−5^, bootstrapping across individuals) **(E)** Alignment of leading principal components of individual variation onto the subspace of sensitivity matrices for four models: the heterogeneous model with uncorrelated noise, the heterogeneous model with uncorrelated noise and network structure in the parameter (Fig. S7A), the heterogeneous model with correlated noise (Fig. S7C), and the heterogeneous model with correlated noise (Fig. 5A) for comparison. Incorporating network structure in the *G* parameter modestly improves fit values but fails to capture PC3 as well as with correlated within-network noise (Fig. 5). Incorporating network structure across E & I pairs (see panel C) yields similar improvements in fit and similar ability to capture PC3 as with network structure only between E-E pairs (see Fig. 5).

**Figure S8.**
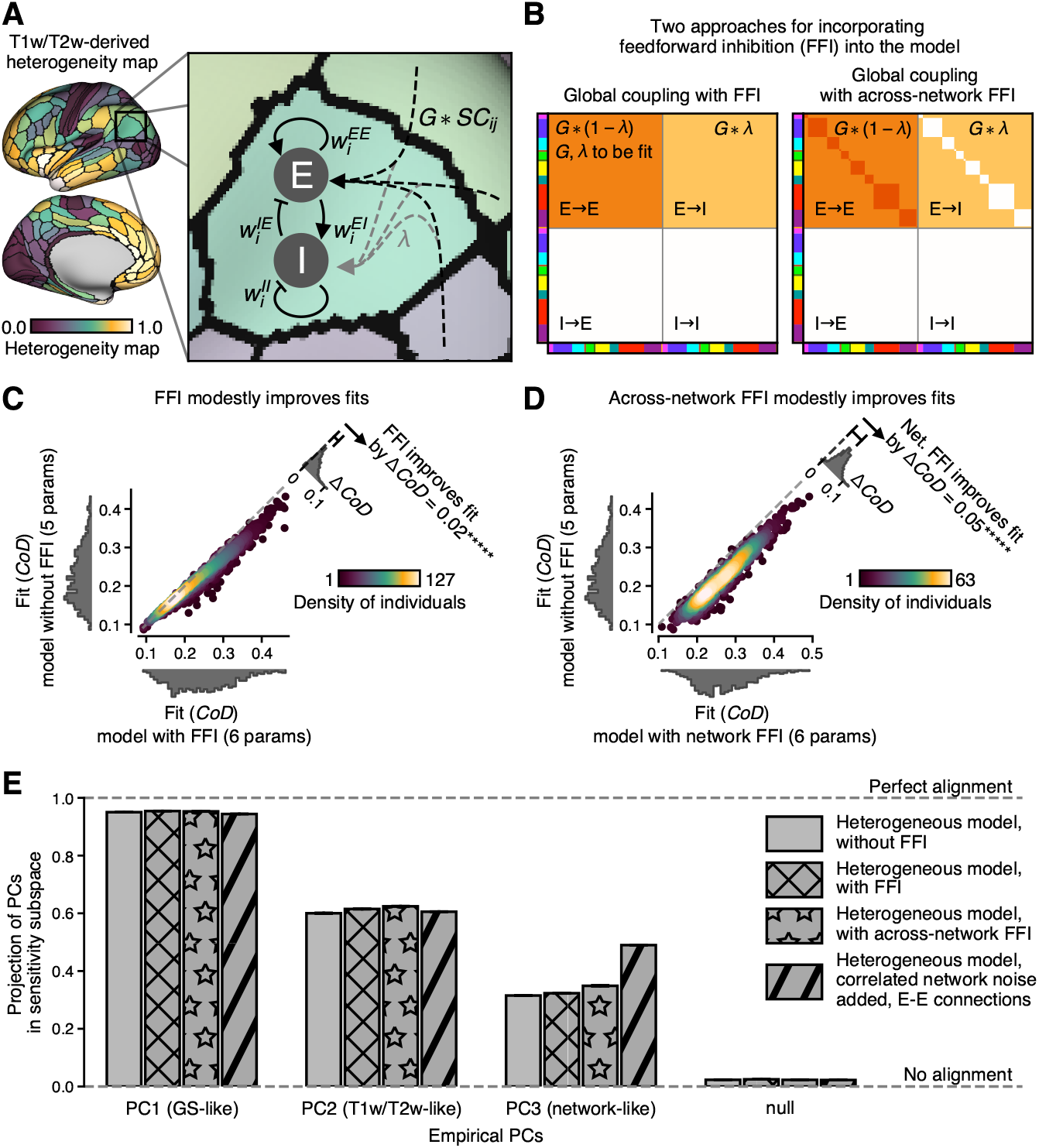
Supplement to Fig. 5 showing an extension of the model incorporating feedforward inhibition (FFI) (Deco et al., 2014). **(A)** Schematic showing how FFI is incorporated into the model. Each excitatory, across-parcel connection (depicted by a dashed line) is split into two components: one exciting the *E* unit and one exciting the *I* unit. **(B)** Schematic depicting two different configurations for FFI. Left: all excitatory, across-parcel connections are assumed to exhibit FFI. Connections are split into two components, with the *E* −*E* component proportional to *G* * (1− *λ*) and the *E* −*I* component proportional to *G* \**λ*. Right: FFI is only assumed to be present between parcels in different networks. This structure was proposed because negative correlations in FC matrices are predominantly found in across-network edges. In either case, connections exhibiting FFI were modeled so as to conserve firing rates, with E → E connections having reduced strength *G* * (1−*λ*) and E →I connections having strength, *G* * *λ*, which sum to *G* as is used in previous models lacking FFI. **(C, D)** Extending the model using FFI according to either approach (global or network structure; see panel B) modestly improves fits relative to the model without FFI. (*****, *p <* 10^−5^, bootstrapping across individuals) **(E)** Models with and without FFI capture the first and second empirical PCs similarly well. The heterogeneous model with correlated network noise (see Fig. 5) has the same dimensionality as the FFI models (6 parameters) and captures the third PC much better than either of the FFI models, suggesting a better ability to capture individual variation with the same level of model complexity.

**Figure S9.**
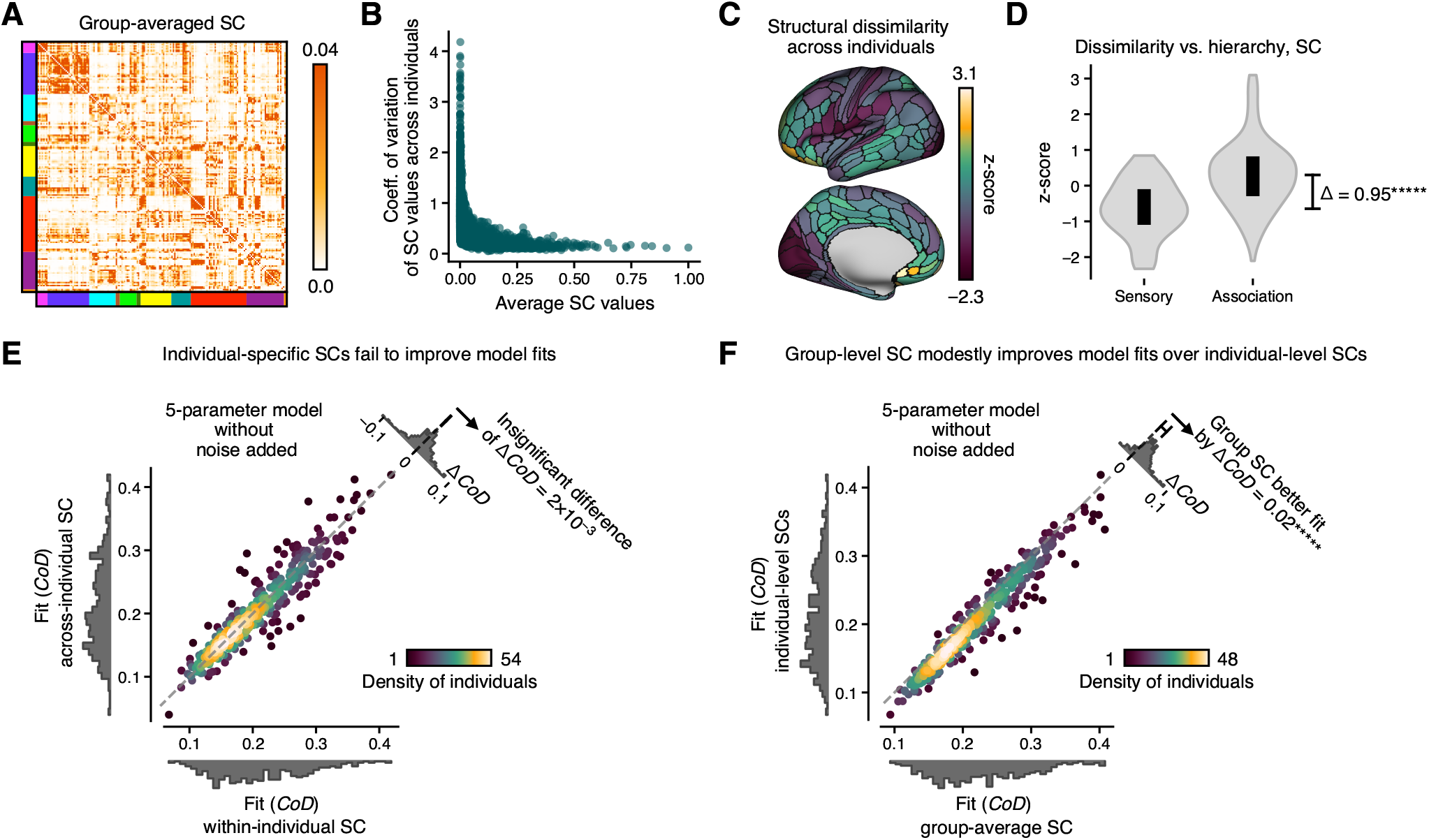
Supplement to Fig. 6. **(A)** Average SC matrix of 328 individuals, which exhibits network structure in sensorimotor regions. This structure largely disappears in the higher-order association regions. **(B)** Coefficient of variation of the subject-level SC matrices across subjects as a function of average SC value. The more strongly connected edges tend to have similar levels of variation, while the most weakly connected edges generally correspond to those with the greatest coefficient of variation. This suggests that variance (whether from individual differences or noise) disproportionately affects the edges with low connectivity (which represents a majority of edges). **(C)** Topography of dissimilarity across subjects for empirical SC matrices. The dissimilarity of parcel is defined as *E* [1 −*r*_Spearman_ (*SC*_*i*_ (*s*_*p*_), (*SC*_*i*_ (*s*_*q*_))], where *E* is the expected value over all pairs of subjects *p* and, *r*_Spearman_ () is the Spearman correlation, and *SC*_*i*_ (*s*_*p*_) is the structural connectivity vector of parcel for subject *s*_*p*_, following the example of Mueller et al. (2013). This shows a pattern similar to the cortical hierarchy map (Fig. 3F), with relatively low dissimilarity within sensorimotor regions and high dissimilarity within association regions. **(D)** Distribution of dissimilarity values by hierarchy, showing disparity between sensorimotor and association regions. Significance was calculated by bootstrapping across cortical parcels. **(E)** Heterogeneous model is insensitive to subject-level information in SC matrices. Given subject-level SC matrices, the model is insensitive to whether the SC matrices are subject-specific or chosen randomly. Subjects are only slightly better fit by the model using their own SC matrix than a randomly chosen subject’s SC matrix. This holds for models with correlated noise (Fig. 6B). Significance was calculated by bootstrapping across subjects. **(F)** Heterogeneous model with and without correlated noise negatively affected by noisiness of individual-level SC matrices. All subjects are fit similarly or better by a model using the group-averaged SC matrix than the subject-level SC matrices. This also holds for models with correlated noise (Fig. 6C). Significance was calculated by bootstrapping across subjects.

**Figure S10.**
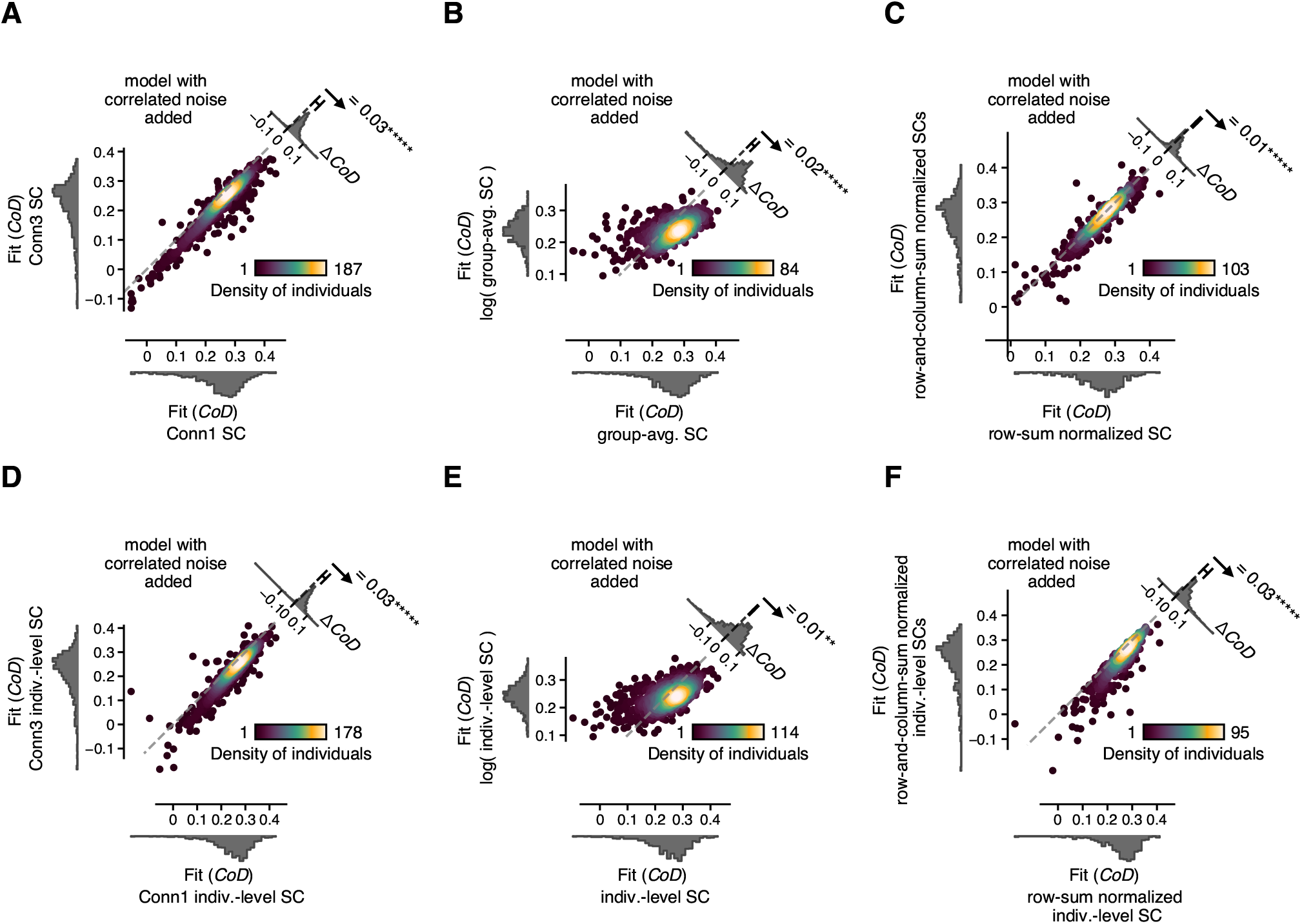
Supplement to Fig. 6. Alternative methods of estimating SC produce similar but generally slightly lower fit values than the estimate of SC used throughout this study. **(A, D)** The Conn1 estimate of SC yields slightly better fit values than Conn3, for both group- and individual-level SC. (*****, *p <* 10^−5^, bootstrapping across individuals) **(B, E)** Taking the logarithm of streamline counts before normalizing generally produces slightly lower fit values, for both group- and individual-level SC. (**, *p <* 10^−2^; *****, *p <* 10^−5^, bootstrapping across individuals) **(C, F)** Normalizing across the entire SC matrix produces slightly lower fit values than normalizing across each row of the SC matrix separately, for both group- and individual-level SC. (*****, *p <* 10^−5^, bootstrapping across individuals).

**Figure S11.**
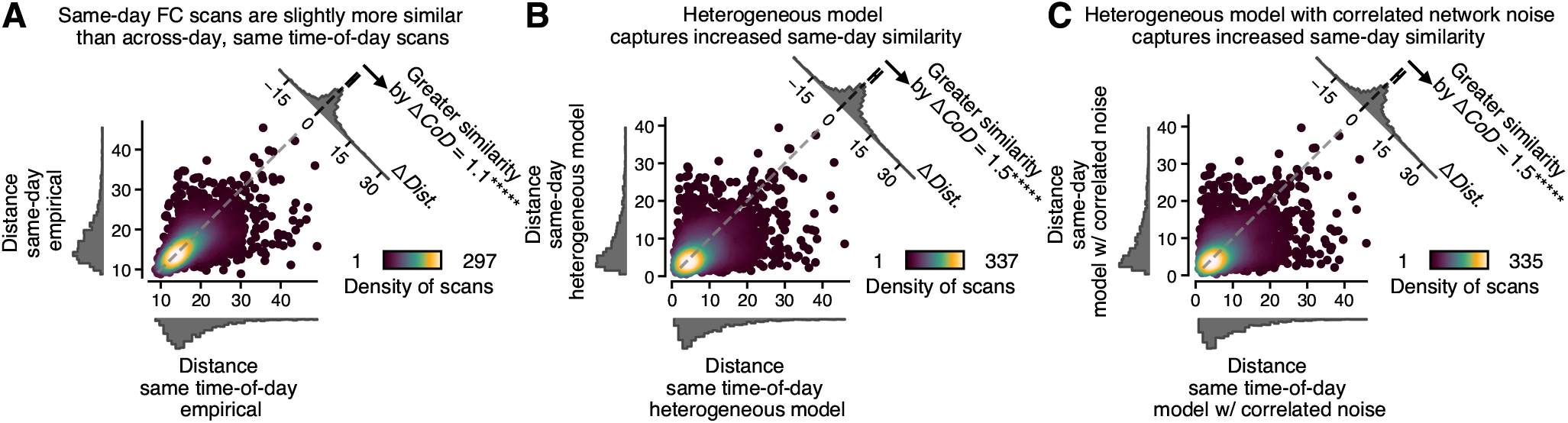
Supplement to Fig. 7. **(A)** Individual-level FC is consistent over time, with same-day scans being slightly more similar than same time-of-day scans taken days apart. Comparing two scans from the same individual, either from the same day or the same time-of-day on a different day, we found that same-day scans are more similar (Δ*Dist*. = 1.1 on average). This difference is small compared to the across-versus within-subject difference (Δ*Dist*. = 6.2 on average, Fig. 7). This suggests that within-individual variation is considerably lower than variation across subjects. (*****, *p <* 10^−5^, bootstrapping across individuals) **(B)** The heterogeneous model captures the increased similarity of same-day scans, as evidenced by having a similar Δ*Dist*. as the empirical data in Panel A. This suggests this model may be adequate for modeling within-individual variation as well, which is a potential future research direction. (*****, *p <* 10^−5^, bootstrapping across individuals) **(C)** Same as Panel B, except for the heterogeneous model with correlated network noise. This extended model also captures the increased same-day similarity. (*****, *p <* 10^−5^, bootstrapping across individuals).

